# Argonaute 1 associates with nucleolar snoRNP machinery and regulates pre-rRNA levels and cell growth

**DOI:** 10.64898/2026.10.19.759891

**Authors:** Rucha Y Hebalkar, Nils Schumann, Andriy S Yatsenko, Rui João Loureiro, Katarina Gresova, Steffen Guenzel, Panagiotis Alexiou, Markus Seiler, Janusz M Bujnicki, Halyna R Shcherbata

## Abstract

Argonaute proteins are best known as cytoplasmic effectors of small-RNA-mediated gene silencing, although their compartment-specific functions remain incompletely understood. Here, we identify a previously unrecognized nucleolar role for Drosophila Argonaute 1 (AGO1). We observed stress-induced redistribution of AGO1 into cytoplasmic and nuclear ribonucleoprotein assemblies, whereas a distinct AGO1 pool consistently localized to Fibrillarin-positive nucleoli. Co-immunoprecipitation further showed that AGO1 associates with Fibrillarin-containing complexes in vivo. Re-analysis of AGO1-associated small-RNA sequencing data revealed snoRNAs, particularly C/D box snoRNAs, among AGO1-bound RNA species and increased snoRNA representation following AGO1 overexpression. Structural modelling demonstrated that simultaneous molecular interaction of AGO1 and the Fibrillarin-containing snoRNP machinery with the same snoRNA is stereochemically feasible, providing a structural framework for their potential association within a nucleolar snoRNP assembly. Functionally, AGO1 depletion reduced pre-rRNA levels and decreased both nucleolar and cell size in the follicular epithelium. Depletion of Dicer-1 did not reproduce these growth defects, arguing against a general consequence of impaired miRNA biogenesis. Together, these findings identify AGO1 as a nucleolar regulator that associates with snoRNP machinery and links small-RNA biology to pre-rRNA homeostasis and cell growth.

**Short summary:** This study identifies *Drosophila* AGO1 as a nucleolar regulator that associates with Fibrillarin-containing snoRNP machinery and links small-RNA biology to pre-rRNA homeostasis and cell growth.

**Key Points:**

- AGO1 is identified as a regulator of nucleolar RNA biology.
- Biochemical, RNA and structural analyses connect AGO1 to snoRNP machinery.
- Nucleolar AGO1 links small-RNA control to pre-rRNA homeostasis and cell growth.

## INTRODUCTION

Sudden environmental changes disrupt cellular homeostasis, requiring cells either to restore the original state or to establish a new adaptive state, depending on stress intensity and duration (1,2). To address these challenges, cells depend on tightly regulated mechanisms of gene expression and protein synthesis. One immediate protective response of cells is to reduce global protein translation before switching to major shifts in gene expression (3). Two key regulatory mechanisms that orchestrate this translational slowdown are miRNA-mediated mRNA silencing and the sequestration of mRNAs into dynamic, membraneless structures called ribonucleoprotein (RNP) granules (4–6). These processes fine-tune the cellular stress response by transiently repressing translation and reorganizing mRNA availability. In parallel, cells also regulate ribosome production itself. Stress can rapidly inhibit early rRNA processing and retain unprocessed rRNA within the nucleolus, thereby coordinating translational shutdown with ribosome biogenesis and preserving nucleolar integrity (7). This places the nucleolus at the center of stress adaptation, linking acute changes in protein synthesis with longer-term control of ribosome production.

miRNA-mediated gene silencing is guided by small non-coding RNAs known as miRNAs, which bind to AGO proteins and serve as sequence-specific guides for silencing target mRNA (Figure 1A). Argonaute (AGO) proteins are central to RNAi pathways, serving as core components of the RNA-induced silencing complex (RISC), which orchestrates post-transcriptional gene silencing via miRNA- or siRNA-guided interactions with target mRNAs (8).

**Figure 1.**
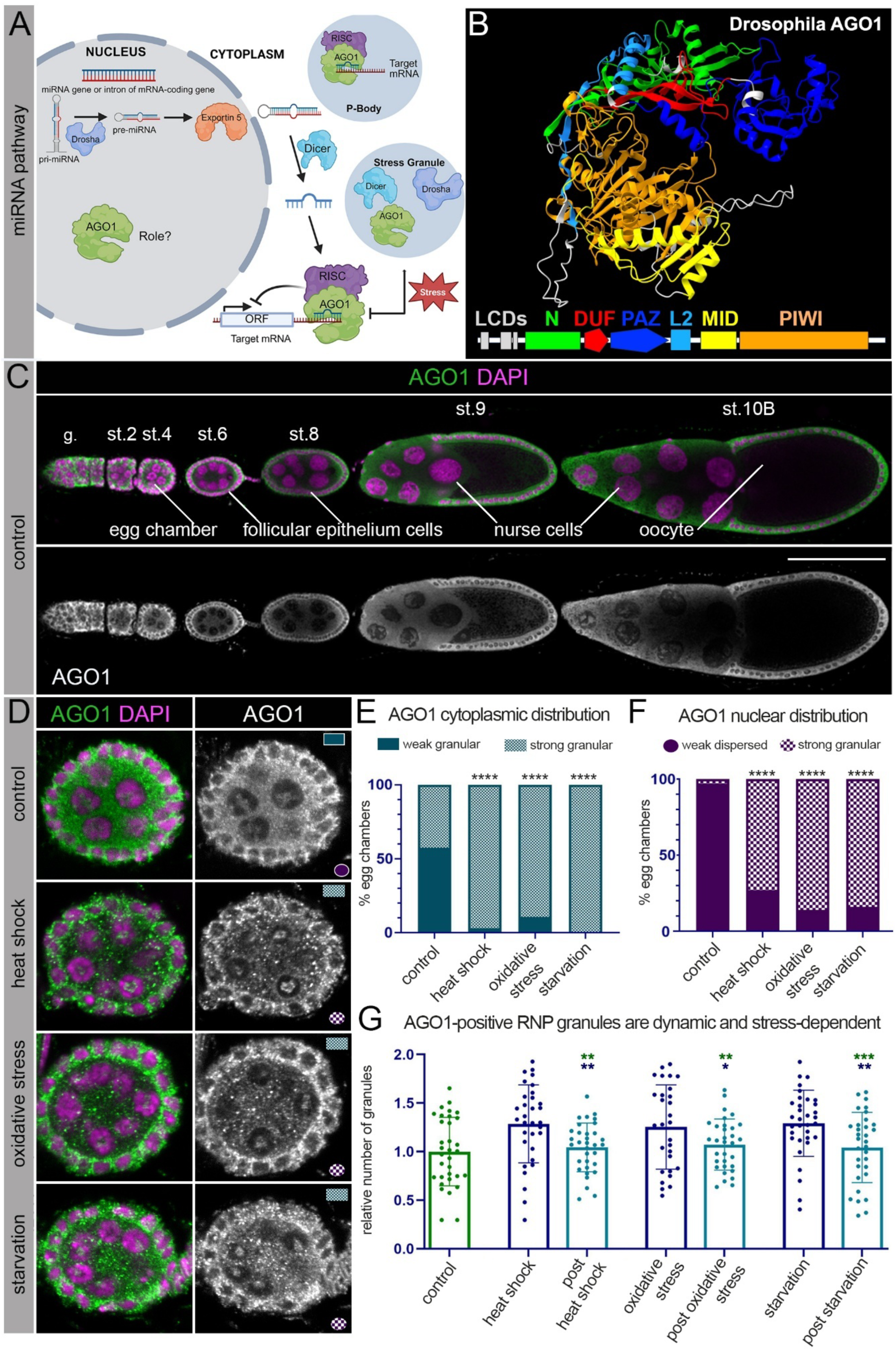
AGO1 accumulates into cytoplasmic and nuclear granules *in vivo* upon stress. **A.** A schematic of the miRNA pathway shows that miRNAs are initially processed in the nucleus by the RNase III enzyme Drosha, which generates precursor miRNAs (pre-miRNAs). These pre-miRNAs are then exported to the cytoplasm, where they are further cleaved by the RNase III enzyme Dicer to produce mature miRNA duplexes. One strand of the duplex is loaded onto an Argonaute (AGO) protein within the RNA-induced silencing complex (RISC), guiding the complex to silence specific target mRNAs. Under cellular stress conditions, components of the miRNA biogenesis pathway, including AGO proteins, relocalize into RNA-protein granules such as processing bodies (PBs) and stress granules (SGs). During stress, cytoplasmic Dicer and the normally nuclear Drosha are also sequestered into SGs, while cytoplasmic AGO1 accumulates in the nucleus. These changes collectively impair miRNA processing and function, suggesting a stress-responsive reorganization of the pathway. **B.** Shows the *Drosophila* AGO1 protein structure as found in the AlphaFold protein structure database developed by Google DeepMind and EMBL-EBI (ID: AF-Q7KY08-F1-v6). The model has a high average pLDDT of 84.94 making it a reliable source for the structure of AGO1. AGO1 contains multiple domains which were identified based on sequence by using SMART analysis. (1) N terminal domain (green): compact α/β fold contributing to guide RNA positioning and conformational stability, (2) Linker region L1 (“DUF1785”, red), (3) PAZ domain (dark blue) interacting with 3′ ends of small RNAs, (4) Linker region L2 (Light blue), (5) MID domain (yellow) interacts with the 5′ phosphate of the guide RNA, (6) PIWI domain (orange) RNase H like fold which in Drosophila AGO1 has no slicer activity. **C.** Shows *Drosophila* ovariole stained with anti-AGO1 (green) and DAPI (magenta) (marks the nuclei). Ovariole made up of germarium (g.) and progressively developing egg chambers from stage 2 - stage 10B (st.2 - st.10B) is shown. Each egg chamber contains germline nurse cells and an oocyte and is encapsulated by somatic follicular epithelium cells. Scale bar = 100µm. **D.** Early-stage egg chambers (st. 2-4) stained with anti-AGO1 (green) and DAPI marking the nuclei (magenta) are shown. In the cytoplasm, the AGO1 distribution pattern is weakly granular and weakly dispersed in the nucleus under control conditions. Upon various stresses, AGO1 accumulates into many more distinct granules in the cytoplasm and the nucleus (strong granular). **E.** Graph shows the percentage of egg chambers with weak granular and strong granular cytoplasmic AGO1 distribution pattern in control and upon various stresses. **F.** Graph shows the percentage of egg chambers with weak dispersed and strong granular nuclear AGO1 distribution pattern in control and upon various stresses. **G.** Graph shows increase in the number of cytoplasmic AGO1 granules upon different stresses and their decrease after stress removal relative to the control. Granules per µm² were counted using Cell Profiler 4.3.1. Wild-type *Oregon-R* flies were used. n≥30 egg chambers analyzed for each condition, three biological replicates. Two-Way Tables and Chi-Square test were applied for statistical analyses in E and F. Student’s t-test was applied for statistical analyses in G. Error bars represent standard error of the mean (SEM). *p≤0.05, **p≤0.01, ***p≤0.001, ****p≤0.0001. Images in D are single Z-sections. Scale bars = 5µm.

In cellular stress responses, miRNAs play a crucial role by either restoring the original expression pattern or establishing a new expression pattern (9,10). At the same time, miRNA biogenesis is extremely sensitive to stress (11,12). Stress triggers the formation and reorganization of membraneless RNP granules (Figure 1A). miRNAs and RISC components, including AGO proteins, have been reported to be localized to RNP granules such as stress granules (SGs) and processing-bodies (PBs) under stress conditions (13). Some RNP granules, including nuage and sponge bodies, are specialized germline condensates involved in post-transcriptional RNA regulation during germ-cell development (14). These RNP condensates are primarily formed through liquid-to-liquid phase separation of RNA-binding proteins that contain low-complexity sequence domains (LCDs), which often are part of intrinsically disordered protein regions (IDRs) lacking defined structure (15,16). Apart from SGs and PBs in the cytoplasm, various types of RNP granules such as nucleoli, Cajal bodies, and paraspeckles are present in the nucleus (5,17,18).

Among these nuclear RNP compartments, the nucleolus is the central site of ribosome biogenesis, where precursor rRNAs are transcribed, chemically modified, processed, and assembled with ribosomal proteins into pre-ribosomal subunits (17,19,20). These maturation steps are guided in part by snoRNAs, which direct site-specific rRNA modifications and contribute to accurate ribosome assembly (21–23). Beyond their canonical role as rRNA modification guides, snoRNAs are increasingly recognized as multifunctional regulators of ribosome production, stress responses, and disease-associated cellular states (24–27). This raises the question of how snoRNAs are selected, stabilized, and positioned within nucleolar RNP complexes, and whether additional RNA-binding proteins contribute to this process (28,29). One class of candidates that could potentially link small-RNA recognition to RNP compartmentalization is the Argonaute family.

Although AGO proteins have been reported to localize to cytoplasmic stress granules and processing bodies, it remains unclear whether they are also recruited to other RNP compartments where they acquire additional regulatory functions. This possibility is supported by growing evidence that AGO proteins act beyond canonical cytoplasmic RNA silencing. For example, fly and human AGO proteins can undergo lipid-mediated phase separation on the endoplasmic reticulum membrane, forming RNP granules that regulate nascent-peptide ubiquitination (30). In Drosophila, AGO1 has also been implicated in transcriptional silencing through transgene co-suppression and heterochromatin formation, as well as in cell-growth control through transcriptional repression of the Myc proto-oncogene (31,32). Similarly, the siRNA-pathway Argonaute AGO2 has been shown to act in the nucleus as a chromatin insulator independently of its RNAi activity and Dcr-2, and to associate with euchromatin to regulate RNA polymerase II-dependent transcriptional elongation (33,34).

Nuclear functions of AGO proteins are also well established in mammalian cells, where they can bind gene promoters and recruit chromatin-modifying factors, leading either to transcriptional activation through enrichment of active chromatin marks or to transcriptional repression through repressive chromatin modifications (35,36). AGO proteins have further been shown to link chromatin silencing with alternative splicing (37). More recently, chimeric eCLIP analyses identified endogenous AGO/small RNA/target RNA complexes not only in the cytoplasm, but also in nuclear and chromatin-associated fractions, reinforcing the concept that AGO-mediated RNA recognition is not restricted to cytoplasmic mRNAs (35). Together, these studies reveal dynamic nuclear roles for AGO proteins in gene-expression regulation. However, whether AGO proteins participate in nucleolar RNA-processing pathways, including snoRNA-guided rRNA modification and maturation, has remained unknown.

In this study, we uncover an unanticipated nucleolar role for *Drosophila* AGO1. We first show that AGO1 accumulates in cytoplasmic and nuclear granules under various stress conditions, and that these granules correspond to known membraneless RNP compartments. Beyond its association with SGs and PBs, AGO1 shows a stress-independent enrichment in the nucleolus, where it colocalizes with Fibrillarin that functions as 2′-O-methyltransferase of small nucleolar RNAs (snoRNAs) within the snoRNPs complex. Guided by this finding, we analyzed AGO-bound small RNA sequencing data (38) and uncovered that AGO1 can indeed bind small nucleolar RNAs (snoRNAs). Our data support the idea that AGO1 can integrate into a hybrid snoRNP-like assembly containing Fibrillarin, SNU13, and box C/D snoRNAs. Computational modeling shows that such a molecular assembly is structurally compatible with a simultaneous binding to the snoRNA 5’-terminus by AGO1 and the 2′-O-methylation of the rRNA target by the Fibrillarin complex. Finally, we show that AGO1 function has physiological consequences: AGO1 loss-of-function or depletion reduces both cell and nucleolar size in the follicular epithelium, whereas depletion of another miRNA-pathway component, Dcr-1, does not. Together, these findings reveal that AGO1 acts beyond its canonical cytoplasmic role, contributing to nucleolar snoRNP function, rRNA processing, and ultimately the regulation of cell growth.

## RESULTS

### AGO1 accumulates in cytoplasmic and nuclear granules in vitro and in vivo

To grow and to survive, cells and organisms must constantly adapt to changing environmental conditions. One key adaptive mechanism involves the formation of ribonucleoprotein (RNP) granules, which are specialized subcellular compartments that regulate gene expression during stress. Stress can alter the localization of critical miRNA pathway components, such as AGO1, thereby influencing the efficiency of miRNA-mediated regulation of stress-responsive genes (Figure 1A). Moreover, relocation of such proteins to different cellular compartments may enable them to acquire new functions. To explore this possibility, we examined the subcellular localization of AGO1 under both normal and stress conditions and subsequently investigated whether it may perform novel functions beyond canonical miRNA activity.

*Drosophila* AGO1 is highly conserved (Supplementary Figure 1) and, like other members of the Argonaute protein family, contains several conserved domains essential for small-RNA-mediated gene regulation. Among these, the Piwi-Argonaute-Zwille (PAZ) domain, which binds the 3′ end of single-stranded guide RNAs, and the P-element-induced wimpy testis (PIWI) domain, which adopts an RNase H-like fold, are central to AGO1 function in the miRNA pathway (Figure 1B). The PAZ domain contains binding pockets that anchor the 3′ and 5′ ends of small RNAs, ensuring proper guide strand positioning (39,40). The PIWI domain is a structural homolog of the DNA-directed RNA endonucleases such as RNase H and contains the catalytic center required for target RNA cleavage. Two further domains, the N-terminal domain and the MID domain assist in the positioning of the PAZ and PIWI domains correctly to interact with small RNAs and also contribute to the formation of the RNA-binding groove (41–43). However, *Drosophila* AGO1 lacks the key residues necessary for this slicer activity, rendering it catalytically inactive (44–46). This observation is important because it suggests that in the miRNA pathway, gene silencing primarily occurs through translational repression and mRNA destabilization, rather than direct target cleavage. This would also allow for more flexible and reversible regulation of gene expression, which is especially crucial during dynamic conditions like cellular stress. In addition, a missing slicer-activity may enable AGO1 to interact with other coding and non-coding RNAs without inducing their degradation.

To study the distribution pattern of AGO1 *in vivo*, we used *Drosophila* oogenesis as a stress-sensitive model (47). *Drosophila* ovaries provide an excellent system to study stress responses because oogenesis is a highly regulated developmental process that is exceptionally sensitive to both physiological and environmental stress. Each ovary is composed of ovarioles, which contain a series of developing egg chambers arranged in a temporal progression (Figure 1C). At the anterior tip of each ovariole lies the germarium, where germline stem cells divide to produce cystoblasts that give rise to 16-cell germline cysts. Within each cyst, one cell becomes the oocyte while the remaining 15 differentiate into nurse cells, which support oocyte development by providing RNAs, proteins, and organelles. The germline cells are surrounded by somatic follicle cells, which encapsulate the cyst to form the egg chamber and contribute essential signals for oocyte maturation. This coordinated interaction between germline and somatic cells makes oogenesis a powerful model to dissect stress-sensitive regulatory pathways, particularly cell growth and differentiation.

To investigate how AGO1 localization in the ovary responds to stress, we subjected flies to several stress conditions, namely heat shock, oxidative stress, and nutrient deprivation. Changes in the distribution of AGO1 were then analyzed using immunohistochemistry. Under normal conditions, the AGO1 localization pattern in the cytoplasm was weakly granular in about 60% of the egg chambers, while it was weakly dispersed in the nucleus (Figure 1D). Upon application of different stresses, AGO1 protein assembled in distinct granules in the cytoplasm and in the nucleus with varying frequency depending on the type of stress applied (Figure 1E-F). In the cytoplasm, 100% of the egg chambers showed AGO1 accumulating in distinct granules upon starvation (Figure 1E). 95% and 90% of the egg chambers showed AGO1 accumulating in distinct granules in the cytoplasm upon heat shock and oxidative stress, respectively (Figure 1E). In the nucleus, AGO1 displayed a strongly granular pattern in 88% of the egg chambers upon oxidative stress, in 85% of the egg chambers upon starvation and in 75% of the egg chambers upon heat shock (Figure 1F). We observed that upon removal of each of the stresses, cytoplasmic AGO1 granules reduced in number, indicating a reversible stress response (Figure 1G). These data show that AGO1 accumulates in cytoplasmic and nuclear granules *in vivo* in *Drosophila* ovaries, which is enhanced by different stresses. Interestingly, AGO1 is predominantly known to function in the cytoplasm to carry out miRNA-mediated gene silencing. Its presence in the nucleus and accumulation in nuclear granules upon stress suggests that it has nuclear functions that might be important for cellular stress response.

### AGO1 associates with various cytoplasmic and nuclear membraneless organelles

To investigate the nature of AGO1 granules and explore novel functions of AGO1, we performed co-immunostaining with AGO1 and well-known ribonucleoprotein (RNP) granule markers to examine if they colocalize in the *Drosophila* germline under stress. We observed colocalization of cytoplasmic AGO1 with various markers for PBs (DCP1, Me31B, Staufen and Yps) and SGs (Hsp70) (Supplementary Figure 2A). This colocalization was enhanced under stress and was mostly observed around the nuclear periphery in a cloud-like pattern (Supplementary Figure 2B). SGs are sites enriched in untranslated or poorly translated mRNAs formed mainly during sub-optimal environmental conditions (5,48,49). AGO proteins interact with molecular chaperones such as Hsp70, which assist in their proper folding, stabilization, and the loading of small RNAs during RISC assembly (50). Under stress conditions, Hsp70 may also help maintain AGO function and facilitate its localization to SGs, linking RNA silencing to the cellular stress response (45,51). PBs contain translationally repressed mRNAs with mRNA decay machinery in the cytoplasm (52). Thus, colocalization of AGO1 with SGs and PBs was not unexpected due to its function in miRNA-associated gene silencing (8,13,53,54).

In contrast, AGO1 colocalization with nuclear RNP granules was unforeseen. Specifically, nuclear AGO1 was found to colocalize with the Cajal body component Coilin and with the polypyrimidine tract-binding protein PTB (also known as Heph) (Supplementary Figure 2A-B). Cajal bodies are sites for assembly and modifications of small nuclear ribonucleoproteins (snRNPs) involved in mRNA splicing (55). PTB is a predominantly nuclear RNA-binding protein that regulates pre-mRNA processing, particularly alternative splicing (56). Intriguingly, the most prominent colocalization of nuclear AGO1 was detected with Fibrillarin, a nucleolar marker, regardless of stress (Supplementary Table 1).

### Nuclear AGO1 colocalizes with Fibrillarin

Under control conditions, we observed a strong colocalization between nuclear AGO1 and Fibrillarin (Figure 2A). Upon heat shock, nuclear AGO1 formed a more compact distribution pattern as compared to control and this was similar to the redistribution pattern of the nucleolus marked by Fibrillarin. Nuclear AGO1 and Fibrillarin maintained their colocalization upon exposure to oxidative stress and starvation. However, in these stress conditions, the overall nuclear distribution of AGO1 remained unchanged compared to the control condition (Figure 2A). We further investigated if the compact distribution pattern of both nuclear AGO1 and Fibrillarin observed upon heat shock would be reversible upon returning flies to control temperature (25°C). It was observed that upon heat shock removal, both nuclear AGO1 and Fibrillarin distribution started to decompress and gradually recover (Figure 2). In addition, under oxidative stress and starvation, cytoplasmic AGO1 distribution did not change compared to the control and remained dispersed throughout the cytoplasm. Upon heat shock, cytoplasmic AGO1 formed clouds around the nucleus as observed previously in Supplementary Figure 2B.

**Figure 2.**
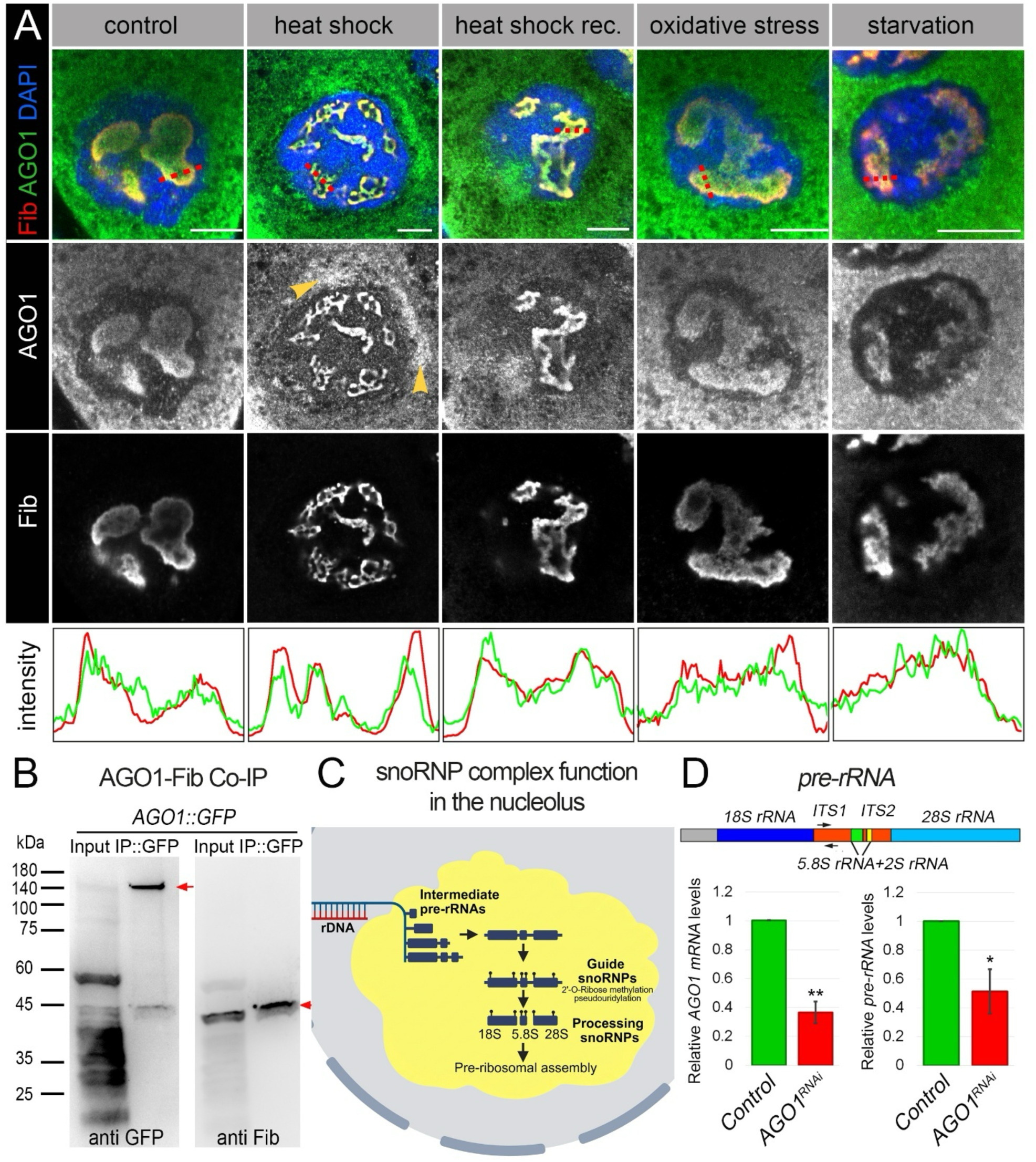
AGO1 colocalizes with the nucleolar marker, Fibrillarin, and it can affect pre-rRNA levels. **A.** Images depict the distribution pattern of AGO1 (green) and nucleolus marked by Fibrillarin (red) under different conditions in late-stage egg chambers (st.8-10). AGO1 strongly colocalizes with Fibrillarin under all conditions (yellow). Upon heat shock, AGO1 redistributes to a more compact structure in the nucleus similar to the redistribution of the nucleolus (Fibrillarin). Note also the perinuclear accumulation of AGO1 in the cytoplasm in response to heat shock (yellow arrows). Upon arsenite treatment and starvation, the AGO1 nucleolar pattern remained unchanged compared to control, but irrespective of the condition, AGO1 colocalized with Fibrillarin. Upon returning flies to control temperature (25°C) post-heat shock, both AGO1 and Fibrillarin start decompressing and restoring to a normal, control-like pattern. Graphs show the intensity profile of AGO1 (green) and Fibrillarin (red) across the red dotted line shown in the merged images. All images are single Z-sections. Scale bars = 10µm. **B.** AGO1 associates with Fibrillarin-containing complexes in Drosophila ovaries. Co-immunoprecipitation (Co-IP) was performed from ovary lysates expressing AGO1::GFP using anti-GFP antibodies to pull down AGO1-containing protein complexes. Western blot analysis of the immunoprecipitated material revealed the presence of endogenous Fibrillarin, detected with an anti-Fibrillarin antibody, demonstrating that AGO1 and Fibrillarin are present in the same protein complex in vivo. Input and IgG control lanes confirm specificity of the pull-down. **C.** Schematic representation of the nucleolus and the function of the snoRNP complex. The nucleolus is a membraneless nuclear structure where rRNA is transcribed, processed, and assembled into ribosomes. Precursor rRNA (pre-rRNA) undergoes extensive chemical modifications, including 2′-O-methylation and pseudouridylation, which are essential for the proper folding, stability, and function of mature rRNA. These modifications are guided by small nucleolar ribonucleoprotein (snoRNP) complexes, which consist of a guide snoRNA and core proteins such as the methyltransferase Fibrillarin (for 2′-O-methylation) and pseudouridine synthases. The guide snoRNA base-pairs with specific sequences in the pre-rRNA, directing the enzymatic activity to defined nucleotide positions. After chemical modifications and processing steps, the mature rRNAs are assembled with ribosomal proteins into functional ribosomal subunits, which are then exported to the cytoplasm to participate in protein synthesis. **D.** Schematic depicts *Drosophila* pre-rRNA transcript which consists of 18S rRNA, 5.8S rRNA, 2S rRNA and 28S rRNA that are separated by internal transcribed spacer (ITS) regions. Black arrows indicate the location of pre-rRNA specific primers used for qPCR. Graphs show the relative mRNA levels of AGO1 and relative pre-rRNA levels upon downregulation of AGO1 in follicular epithelium cell clones (*act > AGO1^RNAi^, GFP^+^*) and upon overexpression of AGO1 in follicular epithelium cell clones (*act > AGO1^OE^, GFP^+^*). CD2 flip-out line (*hsFlp; act>CD2>Gal4, UAS-GFP*) was used as control. Tubulin was used as endogenous control. Relative RNA expression is represented as Mean±SEM, ≥4 biological replicates. Student’s t-test was applied for statistical analyses, *p≤0.05, **p≤0.01.

Within the nucleolus, our immunofluorescence analysis revealed clear colocalization of AGO1 and Fibrillarin. This observation is supported by fluorescence intensity profiles (Figure 2A, lower panel) and 3D reconstruction using IMARIS software, both demonstrating substantial spatial overlap between the two proteins. To determine whether this colocalization is accompanied by a biochemical association, we performed co-immunoprecipitation using AGO1::GFP and an anti-Fibrillarin antibody (Figure 2B). Western blot analysis confirms that AGO1 and Fibrillarin are present in the same protein complex, supporting the existence of a tightly associated interaction within the nucleolar environment.

The nucleolus is the primary site of ribosome biogenesis with its size and shape being highly sensitive to stress conditions. In the nucleolus, ribosome biogenesis is initiated by ribosomal DNA (rDNA) transcription (Figure 2C), producing long precursor rRNAs (pre-rRNAs) that subsequently undergo pre-rRNA modifications (e.g., methylation), extensive endonucleolytic and exonucleolytic cleavage, and conformational folding, followed by the initial stages of ribosomal subunit assembly (57,58). Small nucleolar RNAs (snoRNAs), in complex with specific proteins form small nucleolar ribonucleoprotein (snoRNPs) granules, guide site-specific 2’-O-methylation and pseudouridylation of the pre-rRNA, that are essential modifications for rRNA stability, folding, and function (59). Following these modifications, the processed rRNA is assembled with ribosomal proteins into pre-ribosomal subunits, which are then exported to the cytoplasm for final maturation.

Since Fibrillarin, a key component of the pre-rRNA modification machinery, has been shown to interact with specific sites of rRNA processing (60), it is plausible that AGO1 may also localize to functionally related nucleolar domains involved in rRNA modification and processing. Therefore, we investigated whether changes in AGO1 levels affect pre-rRNA abundance. In eukaryotes, pre-rRNA is transcribed as a single long precursor that includes the 18S, 5.8S, 2S, and 28S rRNAs (Figure 2D). These are separated by internal transcribed spacers (ITS) and are processed through precise cleavage events to produce the mature rRNAs required for ribosome assembly (61). To measure pre-rRNA levels, we designed pre-rRNA-specific primers in the ITS1 region of pre-rRNA of *D. melanogaster* for subsequent qPCR. Since complete loss of AGO1 is lethal, we used the Flp-out system to generate RNAi clones targeting AGO1 in the follicular epithelium. This approach led to approximately 70% reduction in AGO1 levels, as confirmed by RT-qPCR analysis of *AGO1* mRNA (Figure 2D). Importantly, we found that upon AGO1 downregulation (*act>AGO1^RNAi^*, *GFP*, Figure 2D), the levels of pre-rRNA decreased, suggesting that AGO1 influences pre-rRNA levels. Previous studies have already suggested the involvement of key protein components of the RISC pathway, including human AGO2, in pre-rRNA processing (62). Moreover, there is strong evidence that lack of proper modification of pre-rRNA leads to its degradation or defective processing (63,64). Particularly, 2′-O-methylation guided by Fibrillarin is essential for correct folding, stability, and maturation of pre-rRNA (23,65). Since our data suggest that AGO1 interacts with Fibrillarin in the nucleolus and that pre-rRNA levels depend on AGO1, we hypothesized that AGO1 may influence Fibrillarin-associated snoRNP function and thereby contribute to pre-rRNA homeostasis.in the nucleolus.

Fibrillarin is a highly conserved nucleolar protein that functions as an S-adenosyl-L-methionine (SAM)-dependent 2’-O-methyltransferase, whose primary established function is the 2’-O-methylation of RNA. In addition, a limited number of studies have suggested that Fibrillarin can methylate certain proteins (60,66). In the nucleolus, it plays a central role in pre-rRNA processing by catalyzing site-specific 2’-O-methylation of the ribose moieties in ribosomal RNA precursors. The site specificity of this modification is directed by box C/D snoRNAs (23,29).

AGO1, on the other hand, is a central component of the RNA-induced silencing complex and, as described before, is best known for its cytoplasmic role in miRNA-mediated gene regulation (41,42). It binds mature microRNAs and facilitates their interaction with target mRNAs, typically through complementarity between the miRNA seed region (nucleotides 2-8) and the mRNA. This interaction results in translational repression or mRNA degradation, depending on the degree of base pairing and additional contextual features such as the respective RNA structure and protein co-factors (67).

While these two proteins have traditionally been studied in different contexts - Fibrillarin in nucleolar RNA modification and AGO1 in cytoplasmic RNA silencing - our data suggest that they may form a functional association in the nucleolus. To explore this possibility, we aimed to identify non-coding RNAs that could contribute to such a potential interaction between AGO1 and Fibrillarin in the nucleolus. Understanding which RNA species are shared between AGO1 and Fibrillarin could reveal novel regulatory mechanisms that link RNA silencing and RNA modification pathways and provide deeper insight into nuclear RNA metabolism and gene expression control.

### AGO-associated small-RNA data support an association between AGO1 and snoRNAs

Previous studies have suggested that AGO proteins can bind to small RNAs other than miRNAs, such as tRNA-derived small RNAs (tRFs) and rRNA-derived small RNAs (rRFs) (67–70). Therefore, it was hypothesized that AGO1 can bind other small RNAs in a miRNA-pathway-independent manner. To verify this, we analyzed publicly available data (38) of AGO-bound small RNAs in *Drosophila* S2 cells. To capture AGO-bound small RNAs, the authors biochemically purified AGO ribonucleoprotein complexes followed by preparation of a library of bound small RNAs. The raw data from this study was made available at GEO (GSE129600). For this study, we analyzed three biological replicates of samples from *Drosophila* S2 cells (control) and *Drosophila* S2 cells overexpressing AGO1 (*AGO1^OE^*) (sample numbers - GSM3733652, GSM3733653, GSM3733654, GSM3733657, GSM3733658, GSM3733659).

Analysis of normalized reads from control revealed that the vast majority of the AGO-bound small RNAs turned out to be transfer RNAs (tRNAs) (54.97%) followed by miRNAs (32.39%), small nucleolar RNAs (snoRNAs) (8.15%) and small nuclear RNAs (snRNAs) (4.51%) (Supplementary Table 2 and Figure 3A). The detection of AGO-bound snoRNAs was the most intriguing outcome. SnoRNAs are non-coding RNAs mainly present in the nucleolus that play a major role in processing and modification of rRNAs (71). Binding of AGO to snoRNAs was particularly interesting as our data suggested localization of AGO1 in the nucleolus and its probable role in regulating cell growth.

**Figure 3.**
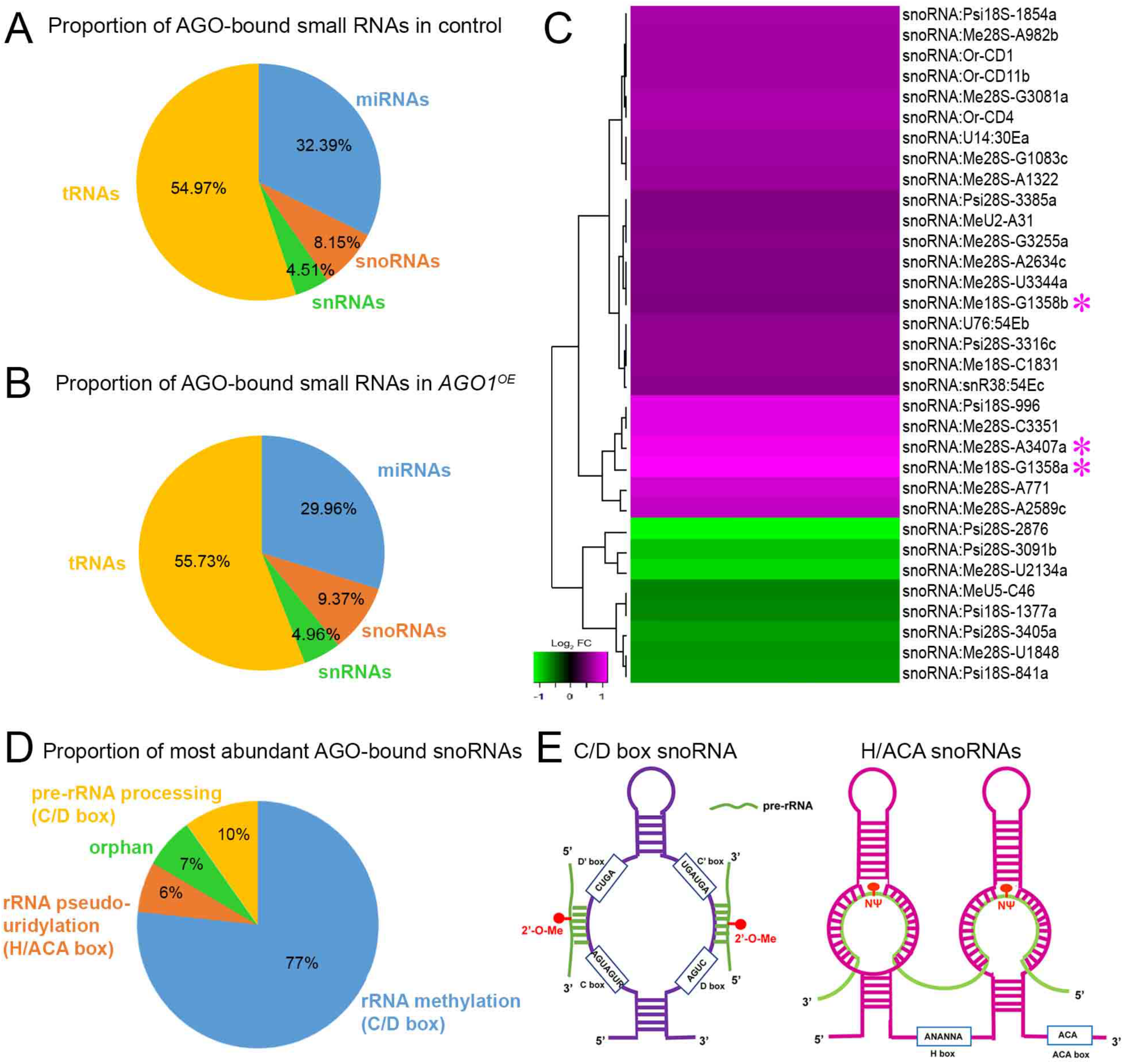
AGO-associated small-RNA datasets reveal enrichment of snoRNAs. **A-B.** Proportion of various AGO-bound small RNAs detected in control (*Drosophila* S2 cells) and in *AGO1^OE^* (*Drosophila* S2 cells overexpressing AGO1) are shown. The majority of small RNAs are tRNAs followed by miRNAs, snoRNAs and snRNAs. There is an increase in the proportion of snoRNAs, snRNAs, tRNAs upon *AGO1^OE^* (see Supplementary Table 2). tRNAs - transfer RNAs; miRNAs - microRNAs; snoRNAs - small nucleolar RNAs; snRNAs - small nuclear RNAs. **C.** Heatmap depicts differentially represented AGO-associated snoRNAs upon AGO1 overexpression (*AGO1^OE^*). 25 out of 33 (∼75%) AGO-associated snoRNAs showed increased representation following AGO1 overexpression (Supplementary Table 4). Analyses in A, B and C was performed using publicly available data (38). **D.** Proportion of the most abundant AGO-bound snoRNAs in *AGO1^OE^*. 22 out of the 30 most abundant snoRNAs in control and *AGO1^OE^* overlapped (see Supplementary Table 3). **E.** Major types of snoRNAs: C/D box snoRNAs guide 2’-O-ribose methylation of pre-rRNAs; H/ACA box snoRNAs guide pseudouridylation of the rRNAs.

Note that the AGO-bound small RNA sequencing data of the control set did not distinguish between AGO1- and AGO2-bound small RNAs (38). Therefore, to increase the chances of discerning AGO1-specific snoRNAs, AGO-bound snoRNA reads from *Drosophila* S2 cells upon AGO1 overexpression (*AGO1^OE^*) were analyzed. The proportion of different AGO-bound small RNAs was similar in *AGO1^OE^*in comparison to control but with a slight increase in snoRNA reads (Supplementary Table 2 and Figure 3B). 174 out of 249 snoRNAs were detected in both control and *AGO1^OE^*.

Supplementary Table 3 lists the 30 most abundant AGO-bound snoRNAs in control set as well as in *AGO1^OE^*. 22 out of the 30 most abundant snoRNAs in control and *AGO1^OE^* overlapped (Supplementary Table 3). We also performed differential expression analysis between *AGO1^OE^* versus control. Out of 33 differentially detected snoRNAs, 25 snoRNAs (∼75%) were 1.5-fold upregulated (Figure 3D and Supplementary Table 4). These results suggest that these 25 snoRNAs were enriched upon AGO1 overexpression.

Based on their conserved motifs and function, snoRNAs may be classified into two main classes (Figure 3E), containing a H/ACA box and a C/D box, respectively (71,72). The H/ACA box snoRNAs display the bipartite consensus motif consisting of the H box (ANANNA) and the ACA box (ACA), and guide the position-specific pseudouridylation of rRNAs (22,71,72). The C/D box snoRNAs are characterized by the presence of two conserved motifs box C (RUGAUGA) and box D (CUGA), and are responsible for 2’-O-ribose methylation of rRNA. The function of some snoRNAs is not known resulting in so-called orphan snoRNAs (73). In our analysis, the majority of abundantly detected AGO-bound snoRNAs were C/D box snoRNAs (Figure 3C–D). C/D box snoRNAs mediate 2′-O-ribose methylation of rRNA and participate in pre-rRNA processing required for mature ribosome formation (74–76). Together, these findings suggest that AGO1 associates not only with miRNAs but also with C/D box snoRNAs, pointing to a previously unrecognized role for AGO1 in nucleolar RNA biology.

### Structural modelling supports the feasibility of an AGO1-associated snoRNP assembly

Our localization, co-immunoprecipitation and small-RNA analyses raised the hypothesis that AGO1 associates with box C/D snoRNP complexes. We therefore used structural modelling to determine whether such an assembly is geometrically feasible. AlphaFold3 (77) did not generate a high-confidence binary AGO1-Fibrillarin complex, prompting us to examine their association within the broader molecular context of snoRNA and the canonical box C/D snoRNP machinery. Given that both proteins function within ribonucleoprotein assemblies, we hypothesized that RNA may facilitate or stabilize their interaction. To test this, we included the sequence of the box C/D snoRNA Me18S-G1359a (CD39b), together with a fragment of its 18S rRNA target encompassing the methylation site. Even with RNA present, AlphaFold3 did not produce a confident model of a ternary AGO1–Fibrillarin–RNA complex with clear biological relevance.

However, AlphaFold3 generated structurally plausible models of each protein in complex with the RNA. In particular, we obtained a model of the complete Drosophila box C/D snoRNP core complex, consisting of Fibrillarin together with its conserved partner proteins (NOP56/58 and SNU13), assembled into the characteristic dimer-of-trimers architecture. In this model, the snoRNA and target RNA adopt a configuration compatible with methylation activity, with the Fibrillarin methyltransferase domain positioned in proximity to the target nucleotide. This result is notable in light of available experimental data. Most experimentally determined structures of eukaryotic Fibrillarin-containing complexes correspond to catalytically inactive conformations associated with RNA processing, in which the methyltransferase domains are rotated away from the substrate RNA. In contrast, catalytically competent conformations have been observed in archaeal box C/D RNP structures, e.g. RSCB record 3PLA, catalytically active substrate-bound box C/D RNP from *Sulfolobus solfataricus*, in which the Fibrillarin homologs are positioned directly over the target nucleotides (78). The inclusion of both guide snoRNA and target RNA in our modeling has apparently prompted AlphaFold3 toward modeling a conformation closely resembling these catalytically active archaeal box C/D RNP structures. In parallel, we obtained a model of Drosophila AGO1 bound to the 5′ terminus of the snoRNA. This is consistent with the well-established preference of AGO proteins for anchoring RNA 5′ ends within their MID domain, a key feature of their interaction with guide RNAs in RNA silencing pathways.

To construct a joint model, we performed protein–protein docking for AGO1 and Fibrillarin using first HDOCK (79) and then HADDOCK (80). The resulting docking results were filtered based on a geometric criterion requiring that the 5′ end of the snoRNA bound to AGO1 could be connected to the remainder of the snoRNA within the Fibrillarin complex without introducing excessive strain or unrealistic distortions. The AGO1–RNA partial model and the Fibrillarin snoRNP partial model were then combined based on the predicted AGO1-Fibrilllarin interface The RNA segment linking the AGO1-bound and Fibrillarin-bound regions was subsequently refined using SimRNA (81). The full assembly was then subjected to all-atom optimization comprising an energy minimization using the GROMACS 2021.2 software suite, with the AMBER99SB-ILDN and AMBER94 force fields, for the protein and the RNA components of the complex, respectively. The minimization removed steric clashes and improved local geometry at protein–protein and protein–RNA interfaces.

The resulting model suggests that AGO1 can interact with the methyltransferase domain of Fibrillarin while simultaneously engaging the snoRNA and the target RNA. In this model, the Fibrillarin complex retains its dimer-of-trimers organization, in which both methyltransferase domains are, in principle, capable of accommodating target RNAs. However, in the case of CD39b, which guides methylation at a single site, only one half of the complex is functionally engaged. In our model, the target RNA pairs with the guide region of the snoRNA and is positioned within the active site of the same Fibrillarin subunit that interacts with AGO1. The SNU13 protein binds the canonical k-turn motif located in the central region of the snoRNA, consistent with established structural data. Notably, binding of AGO1 to the 5′ end of the snoRNA imposes a structural rearrangement that disrupts base pairing between the 5′ and 3′ terminal regions of the snoRNA. As a consequence, all key functional interactions in the model are localized within a single trimeric half of the Fibrillarin complex.

This model is inherently speculative and should be interpreted with caution. Its primary purpose is to demonstrate that a simultaneous interaction of AGO1 and Fibrillarin with the same snoRNA molecule is structurally feasible, without major steric conflicts. The precise molecular details of the AGO1-Fibrillarin interface remain to be determined experimentally. A plausible mechanistic hypothesis is that AGO1 initially binds the snoRNA stem, as suggested by the AlphaFold3 model of the AGO1–snoRNA stem complex. This interaction may enable AGO1 to unwind the snoRNA duplex, thereby facilitating the binding of rRNA to its complementary region within the snoRNA. The complete snoRNP would then assemble. In the last stage, AGO1 leaves the complex and the methylation of the rRNA target by the Fibrillarin domain occurs (Figure 4). Until further data become available, the model provides a working hypothesis that can guide further biochemical and structural studies. In addition, the separate models of AGO1 and Fibrillarin complex with the RNAs can guide the experimental analyses of the Fibrillarin and AGO1 in Drosophila with their RNA binders.

**Figure 4.**
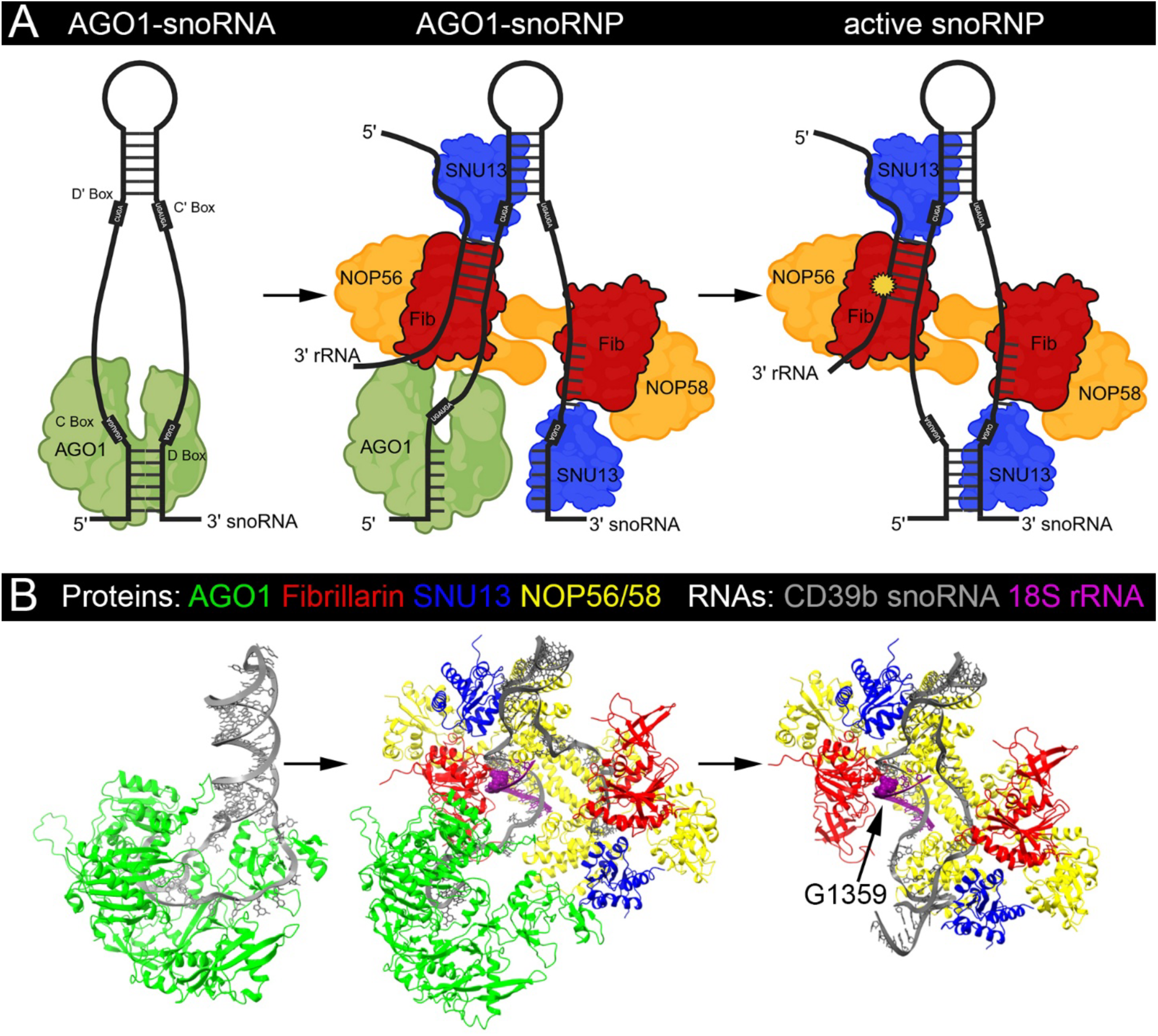
Structural feasibility and working model of an AGO1-associated snoRNP assembly. **A.** SnoRNAs canonically act together with Fibrillarin (red), SNU13 (blue), and NOP56/58 (yellow) in snoRNP complexes to facilitate rRNA methylation. Based on our previous findings, we hypothesize AGO1 (green) is involved in this process, as it directly binds snoRNAs and interacts with Fibrillarin. We propose that AGO1 first binds to the double stranded snoRNA stem, afterwards the snoRNP complex is assembled allowing interactions between Fibrillarin and AGO1 to occur. The presence of AGO1 in the snoRNP complex is proposed to facilitate easier binding of the rRNA to its complementary region on the snoRNA, owing to AGO1’s ability to separate RNA duplexes which could ease tensions occurring during annealing of the rRNA and snoRNA. Once AGO1 leaves the complex and the snoRNA stem structure is restored, methylation by Fibrillarin can occur (yellow star). **B.** Alphafold3 predictions of AGO1 binding to the CD39b snoRNA consistently produced high confidence models (iPTM = 0.84, pTM = 0.84) placing the 5’ end of the RNA in the binding groove of AGO1, supporting AGO1’s ability to bind to the double stranded snoRNA stem. To investigate AGO1’s role within the snoRNP complex, additional models were generated in two steps: first, AlphaFold3 models of the core snoRNP complex (Fibrillarin, SNU13, NOP56/58, CD39b and a fragment of 18S rRNA) were produced; then, protein docking using HADDOCK was performed to generate the full complex. This approach reproduced the proposed open conformation of the AGO1-bound snoRNA while preserving a snoRNP architecture similar to catalytically active complexes observed in archaea (PDB: 3PLA). The snoRNA stem re-anneals once AGO1 is released from the complex, restoring the canonical snoRNP structure and enabling methylation of G1359 (black arrow) in the 18S rRNA. See also Supplementary models.

### AGO1 loss and downregulation affect cell and nucleolar size

Since we propose that, beyond AGO1’s well-established role in post-transcriptional silencing, it may also influence snoRNA-rRNA interactions and facilitate 2′-O-methylation by Fibrillarin, we next aimed to investigate the biological significance of AGO1’s association with the snoRNP complex in the nucleolus. Given that the nucleolus is recognized as the “ribosome factory” and is critical in regulating cell growth, we hypothesized that AGO1 might function within the nucleolus to influence this process. To test this, we needed a model to explore the correlation between nucleolar size and cell growth. Myc, a well-known regulator of cell growth, influences various aspects of ribosome biogenesis, including rRNA transcription, processing, and ribosomal protein production (82). Therefore, we chose Myc as a model to assess potential differences in cell and nucleolar size within the somatic follicular epithelium cells of the *Drosophila* ovary. These cells undergo endoreplication, a process associated with evident cell growth (83). For our study, we induced clones in the follicular cell epithelium using the Flp-out method, such that the same egg chamber would contain control cells and clonal cells with dysregulated Myc expression. It was observed that the clonal cells with downregulated Myc levels (*act>Myc^RNAi^, GFP*) were significantly smaller than neighbouring control cells (Supplementary Figure 3A-B). Moreover, the nucleolar size was significantly reduced in Myc-downregulated clonal cells as compared to control cells (Supplementary Figure 3A-B). Conversely, when Myc was overexpressed (*act>Myc^OE^, GFP*), both the cell size and nucleolar size significantly increased compared to neighbouring control cells (Supplementary Figure 3C-D). Importantly, there was a correlation between cell size and nucleolar size. In summary, the follicular epithelium cell clonal system proved to be a good read-out model to observe differences in the cell and nucleolar size. Therefore, we decided to utilize the system of inducing follicular epithelium cell clones to investigate the potential impact of AGO1 loss and downregulation on cell and nucleolar size.

Initially, we induced AGO1 loss-of-function follicular epithelium cell clones using heat shock inducible FLP/FRT system. GFP-negative AGO1 loss-of-function clonal cells (*hsFlp; FRT42B AGO1^LOF^*) were significantly smaller compared to neighbouring control cells (Figure 5A-B). Furthermore, the nucleolar area was significantly reduced upon AGO1 loss-of-function compared to the control (Figure 5A-B). Subsequently, AGO1-downregulated follicular epithelium cell clones were induced using the Flp-out method. Similar to AGO1 loss-of-function, AGO1 downregulated clonal cells (*act>AGO1^RNAi^*, GFP) were significantly smaller and had reduced nucleolar area compared to neighbouring control cells (Figure 5C-D). Overall, these results suggested that AGO1 loss and downregulation lead to decreased cell and nucleolar size.

**Figure 5.**
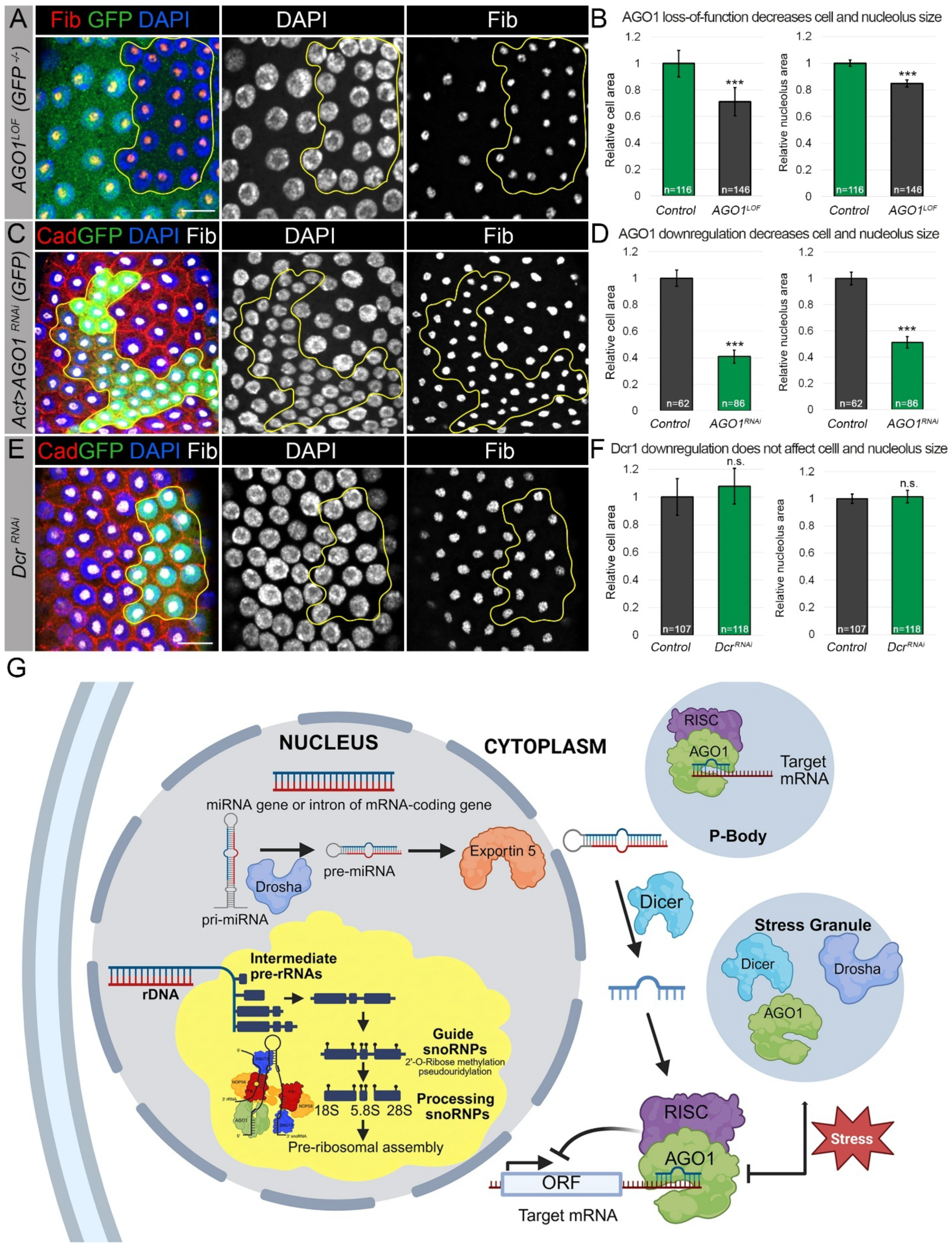
AGO1 expression levels regulate cell and nucleolar size. **A-D. AGO1 loss and downregulation lead to decreased cell and nucleolar size** **A.** AGO1 loss-of-function follicular epithelium cell clones (*hsFlp; FRT42B AGO1^LOF^*, GFP-negative, outlined by yellow line) have smaller cells when compared to the neighbouring heterozygous control cells (*hsFlp; FRT42B GFP/FRT42B AGO1^LOF^*, GFP-positive). DAPI marks the nuclei (blue). Fibrillarin (Fib) marks the nucleoli (red). Fibrillarin staining shows that *AGO1^LOF^* has smaller nucleoli compared to the neighbouring control cells. Images are maximum intensity projections of multiple Z-sections. Scale bar = 10µm. **B.** Mean cell and nucleolar area are significantly decreased in *AGO1^LOF^* clones compared to the respective means of the neighbouring control cells (n=11 clones). Cell and nucleolar area were measured using ImageJ. Mean cell and mean nucleolar areas (Mean±SEM) are represented relative to the control. Statistical significance was assessed using a paired Student’s t-test; ***p≤0.001. **C.** AGO1 downregulated follicular epithelium cell clones (*act>AGO1^RNAi^, GFP,* outlined by yellow line) have smaller cells compared to neighbouring control cells. DAPI marks the nuclei (blue). Fibrillarin (Fib) marks the nucleoli (grey). Cadherin (Cad) marks the cell boundaries (red). Fibrillarin staining shows that act>*AGO1^RNAi^*has smaller nucleoli compared to the neighbouring control cells. Images are maximum intensity projections of multiple Z-sections. Scale bar = 10µm. **D.** Mean cell and nucleolar area are significantly decreased in *AGO1^RNAi^* clones compared to the respective means of the neighbouring control cells (n=10 clones). Cell and nucleolar area were measured using ImageJ. Mean cell area and mean nucleolar (Mean±SEM) are represented relative to the control. Statistical significance was assessed using a paired Student’s t-test; n.s., not significant; ***p≤0.001. **E-F. Dcr-1 downregulation does not affect cell and nucleolar size** **E.** Cells in Dcr-1 downregulated follicular epithelium cell clone (*act>Dcr-1^RNAi^, GFP*) do not differ in size compared to the neighbouring control cells (outlined by yellow line). Fibrillarin staining shows that nucleoli in *Dcr-1^RNAi^* do not differ in size compared to the nucleoli in the neighbouring control cells. DAPI marks the nuclei (blue), Fibrillarin (Fib) marks the nucleoli (grey) and Cadherin (Cad) marks the cell boundaries (red). All images are maximum intensity projections of multiple Z-sections. Scale bars = 10µm. **F.** Mean cell and nucleolar areas did not differ significantly between *Dcr-1^RNAi^* cells and neighbouring control cells (n = 8 clones). Cell and nucleolar areas were measured using ImageJ and are presented relative to the corresponding control values as mean ± SEM. Statistical significance was assessed using a paired Student’s *t*-test; n.s., not significant. **G.** A model that proposes that AGO1 is an RNA-binding protein with distinct functions determined by its subcellular localization and the class of interacting small RNAs. In the cytoplasm, AGO1 binds to miRNAs to mediate silencing of target mRNAs. In the nucleus, AGO1 associates with snoRNAs and may contribute to pre-rRNA homeostasis.

AGO1 loss and downregulation can potentially impact many cellular processes due to its miRNA-associated role in gene silencing. Therefore, the possibility that the cell growth phenotype might be due to the general miRNA-dependent function of AGO1 could not be rejected. To distinguish whether this is miRNA pathway-dependent or -independent phenotype, we sought to examine the impact of downregulating another key component of the miRNA pathway, Dicer-1 (Dcr-1), on follicular epithelium cell size. We induced Dcr-1-downregulated follicular epithelium cell clones using the Flp-out method. Dcr-1 downregulated clonal cells (*act>Dcr-1^RNAi^, GFP*) did not differ in size compared to neighbouring control cells (Figure 5E-F). We also observed that the size of the nucleoli in the Dcr-1 downregulated clonal cells did not show significant difference compared to neighboring control cells (Figure 5E-F). This result argues against the phenotype being a general consequence of impaired miRNA biogenesis and supports a function of AGO1 that is not shared by Dcr-1.

## DISCUSSION

Our study identifies a non-canonical nucleolar role for Drosophila AGO1 in ribosomal RNA metabolism, nucleolar organization, and cellular growth. We propose that AGO1 acts as a compartmentally flexible RNA-binding protein that connects small-RNA pathways with ribosome biogenesis. In this model, AGO1 uses distinct RNA partners in different subcellular environments: miRNAs in the cytoplasm to regulate mRNA stability and translation, and snoRNAs in the nucleolus to influence rRNA processing, nucleolar architecture, and growth. Thus, AGO1 may function not only as a post-transcriptional silencing factor, but also as a broader organizer of RNA-dependent gene-expression control.

Our data show that AGO1 redistributes in cytoplasmic and nuclear granules in a stress-dependent manner. AGO1-positive granules colocalize with several RNP compartments in the Drosophila germline, including stress granules, processing bodies, germline-specific processing bodies, Cajal bodies, and the nucleolus. Localization to stress granules and processing bodies is consistent with the established role of AGO1 in miRNA-mediated translational repression and mRNA turnover. In contrast, its accumulation in nuclear RNP granules, and particularly its prominent association with the nucleolus, suggests that stress redirects AGO1 toward additional RNA-regulatory functions beyond canonical cytoplasmic silencing.

This hypothesis is consistent with accumulating evidence that core small-RNA pathway factors can function outside their canonical compartments. AGO proteins are present in the nucleus and have been implicated in transcriptional gene silencing, chromatin regulation, and alternative splicing (31,32,36,37,84). Other key miRNA biogenesis proteins, including Drosha and Dicer, have also been detected outside their expected compartments (85–87). Alteration of subcellular localization of Drosha was observed in response to stress (88). in addition, alternative splicing produces Drosha isoforms with distinct localizations, including cytoplasmic forms that can still process pri-miRNAs in a DGCR8-dependent manner (85,87). Dicer also has been detected in the nucleus, where it participates in processing double-stranded RNAs that may guide AGO-mediated heterochromatin formation (86). Thus, small-RNA pathway proteins can acquire additional activities depending on their localization, molecular partners, and cellular context (1).

Our study extends this principle by placing AGO1 within the nucleolus, a nuclear RNP condensate where rRNA transcription, modification, processing, and early ribosome assembly are coordinated. The nucleolus is increasingly recognized as a stress-responsive organelle that couples ribosome production to cellular growth and survival (89–91). In this context, AGO1 recruitment to Fibrillarin-positive nucleolar domains may provide a mechanism by which stress-sensitive small-RNA pathways communicate with ribosome biogenesis. Notably, AGO1 colocalizes with Fibrillarin under normal conditions, whereas stress promotes a more compact nucleolar distribution of both proteins. This suggests that AGO1 may have a constitutive nucleolar function that becomes integrated into stress-induced remodeling of nuclear RNP compartments. The previously reported interaction between human AGO2 and Fibrillarin detected by mass spectrometry further suggests that associations between Argonaute proteins and nucleolar components may be evolutionarily conserved (92).

A central concept emerging from our work is that AGO1 may bridge two major RNA-based regulatory systems: cytoplasmic miRNA-mediated translational control and nucleolar snoRNA-guided rRNA maturation. AGO1 colocalizes and physically associates with Fibrillarin-containing complexes, influences pre-rRNA levels, binds non-canonical classes of small RNAs including snoRNAs, and regulates both nucleolar and cell size. These observations suggest that AGO1 contributes to nucleolar RNA-processing pathways in addition to its canonical miRNA-associated function. They also raise the possibility that AGO1 coordinates translational output at two levels: by repressing specific mRNAs in the cytoplasm and by modulating ribosome biogenesis or ribosome quality in the nucleolus.

Mechanistically, AGO1 could influence nucleolar RNA metabolism in several, non-mutually exclusive ways. First, AGO1 may stabilize or retain specific C/D box snoRNAs within nucleolar RNP complexes. Second, analogous to its role in miRNA-guided target recognition, AGO1 may facilitate or stabilize interactions between snoRNAs and pre-rRNA. Third, AGO1 may affect the assembly, positioning, or residence time of snoRNP complexes within Fibrillarin-positive nucleolar subdomains. These possibilities are consistent with our observation that AGO1 interacts with Fibrillarin, associates with snoRNAs, and affects pre-rRNA abundance, nucleolar size, and cell growth.

Our structural and computational analyses support the feasibility of an AGO1-associated snoRNP-like model. Although the full AGO1–Fibrillarin–snoRNA–rRNA complex remains speculative and requires experimental validation, separate modeling of AGO1–RNA and Fibrillarin-containing snoRNP assemblies suggests that AGO1 can bind the 5’ termini of snoRNA and snoRNA-like molecules in a manner compatible with its known guide-RNA-binding mode. These findings suggest that AGO1 may not simply bind snoRNAs passively, but could influence the selection, stabilization, or efficiency of snoRNA–pre-rRNA interactions.

An important implication of this model is that AGO1 may contribute to the plasticity of rRNA modification. Fibrillarin is a major component of C/D box snoRNP complexes and catalyzes 2′-O-methylation of pre-rRNAs, a modification important for rRNA folding, processing, stability, and ribosome function (66). Loss-of-function of C/D box snoRNP components, including Fibrillarin, results in decreased nucleolar size (93). Moreover, nucleolar size is directly associated with RNA polymerase I-dependent rDNA transcription, protein synthesis capacity, and cell growth. By associating with C/D box snoRNAs and Fibrillarin, AGO1 could provide an additional layer of specificity or regulation to snoRNA-guided rRNA modification and processing. This would place AGO1 upstream not only of mRNA silencing, but also of ribosome production and potentially ribosome specialization.

Functionally, AGO1 loss-of-function or downregulation reduces both nucleolar and cell size in the follicular epithelium. Importantly, downregulation of Dcr-1, another key component of the canonical miRNA pathway, did not produce the same phenotype. Thus, the effect of AGO1 on nucleolar and cell growth is unlikely to be a general consequence of impaired miRNA biogenesis. Instead, AGO1 appears to have a separable nucleolar function, consistent with its interaction with snoRNAs and Fibrillarin and its effect on pre-rRNA abundance. In this view, AGO1 acts not only as an effector of mature miRNAs, but also as a regulator of nucleolar capacity that couples small-RNA biology to cellular growth control.

Our findings also refine previous work linking AGO1 to Myc-dependent cell-growth regulation. In Drosophila wing imaginal discs, AGO1 was shown to act as a tumor suppressor by repressing transcription of the Myc proto-oncogene. In that context, AGO1 is enriched at the Myc promoter, interacts with RNA polymerase II and Psi, and AGO1 depletion increases Myc expression, nucleolar size, and cell growth (32). In contrast, we find that AGO1 loss-of-function or knockdown in follicular epithelial cells reduces nucleolar and cell size. Rather than representing conflicting outcomes, these observations suggest that AGO1 engages distinct nuclear growth-control programs depending on developmental and cellular context. In mitotically proliferating imaginal discs, AGO1 may primarily restrain growth through transcriptional repression of Myc. In endoreplicating, differentiating follicle cells, AGO1 may instead support growth by promoting nucleolar rRNA processing and snoRNP-associated ribosome biogenesis. This context dependency strengthens the idea that AGO1 function is determined by cellular state, available RNA partners, and subnuclear compartmentalization.

Our analysis of AGO-bound small RNAs revealed that tRNA-derived reads represent the major fraction of AGO-associated small RNAs. This was expected, because tRNAs are essential for translation and represent a large proportion of total cellular RNA, estimated at approximately 10–15%, compared with substantially lower proportions of miRNAs, snoRNAs, and snRNAs (94). In addition, tRNA-derived fragments are loaded onto AGO proteins and can repress target RNAs through RNAi-like mechanisms (68–70,95). tRFs are also detected in the nucleus, where AGO-loaded tRFs have been proposed to regulate transcriptional silencing and alternative splicing (68,69). In Drosophila, tRFs can suppress translation globally by targeting components of the translation machinery, resembling miRNA-like regulatory mechanisms (70,96). Although our study focuses on snoRNAs because of AGO1’s nucleolar localization and interaction with Fibrillarin, the abundance of AGO-bound tRNA-derived fragments reinforces the broader conclusion that AGO1 can associate with diverse small-RNA classes beyond canonical miRNAs. Different AGO1-bound small RNAs may therefore define distinct functional AGO1 pools across subcellular compartments.

Several questions remain open. First, AGO1 CLIP or RIP-seq experiments in ovaries will be required to define the endogenous AGO1-bound snoRNA repertoire in the relevant tissue context. Second, direct mapping of rRNA 2′-O-methylation, for example by RiboMeth-seq or related approaches, will be necessary to determine whether AGO1 depletion changes modification at specific snoRNA-guided sites. Third, biochemical reconstitution, proximity labeling, or crosslinking-mass-spectrometry approaches will be needed to determine whether AGO1 forms a stable snoRNP-associated complex or interacts transiently with snoRNP components. Finally, rescue experiments using AGO1 variants defective in small-RNA binding, nuclear localization, or disordered regions would help separate its canonical cytoplasmic miRNA function from its nucleolar role.

The involvement of AGO1 in nucleolar growth control has potential implications for diseases associated with abnormal ribosome biogenesis and nucleolar dysfunction. Nucleolar dysfunction is increasingly recognized as a contributing factor in cancer, neurodegenerative disorders such as Alzheimer’s disease, Parkinson’s disease, and amyotrophic lateral sclerosis, and genetic syndromes including Hutchinson–Gilford progeria, Diamond-Blackfan anemia, and Treacher Collins syndrome (89). In autoimmune diseases such as systemic lupus erythematosus, autoantibodies against nucleolar components further underscore the disease relevance of nucleolar pathways (90). Given AGO1’s newly identified role in nucleolar RNA metabolism, AGO-dependent regulation of ribosome biogenesis may represent a conserved disease-relevant mechanism. More broadly, our work suggests that small-RNA effector proteins may regulate not only transcript availability, but also the ribosomal machinery that determines translational capacity.

In conclusion, our study identifies AGO1 as a non-canonical nucleolar regulator that connects small-RNA biology with rRNA processing, nucleolar organization, and cell growth. We propose that AGO1 integrates cytoplasmic microRNA silencing with nucleolar snoRNA-guided rRNA modification and processing, thereby coordinating ribosome biogenesis, translational capacity, and growth, particularly under stress. These findings expand the concept of Argonaute function beyond cytoplasmic mRNA silencing and nuclear chromatin-associated regulation, placing AGO1 within the machinery that controls ribosome production and potentially ribosome specialization. More broadly, our work suggests that Argonaute proteins can act as compartment-specific RNA regulators whose functions are determined by the small RNAs they bind, the protein complexes they engage, and the RNP condensates in which they operate.

## DATA AVAILABILITY

The AGO-associated small-RNA sequencing data analysed in this study are publicly available in the Gene Expression Omnibus under accession number GSE129600. All other data supporting the findings of this study are included in the article and its Supplementary Data. The corresponding model coordinates and analysis scripts will be deposited in a public repository upon acceptance of the manuscript.

## ACKNOWLEDGEMENTS

We thank Mikiko Siomi, Anne Ephrussi, Julius Brennecke and Richard Carthew for providing fly stocks and reagents. We also thank the members of the Shcherbata laboratory for critical reading of the manuscript and helpful suggestions.

## AUTHOR CONTRIBUTIONS

R.Y.H. and N.S. (Formal analysis, Investigation, Methodology, Validation, Visualization, Writing-original draft); R.J.L., K.G. and P.A. (Formal analysis, Methodology, Software, Visualization); S.G. (Formal analysis, Methodology); A.S.Y. (Formal analysis, Methodology, Visualization); M.S. (Conceptualization, Formal analysis, Investigation, Methodology, Validation, Visualization, Writing-review and editing); J.M.B. (Conceptualization, Formal analysis, Investigation, Validation, Supervision, Visualization, Writing-review and editing); H.R.S. (Conceptualization, Formal analysis, Funding acquisition, Investigation, Methodology, Project administration, Resources, Supervision, Validation, Visualization, Writing-original draft, Writing-review and editing).

## Materials and Methods

### Fly stock maintenance

Fly stocks were maintained in vials containing cornmeal-agar prepared with 6.25g/L agar (Serva), 18g/L dry yeast (Saf-Instant), 80g/L corn flour (Zieler & Co.), 22g/L beet syrup (FerdinandKreutzer Sabamühle GmbH), 80g/L malt (Ulmer Spatz), 0.625% propionic acid (Merck), 1.5 g/Lmethylparaben (Sigma). Flies were kept in a controlled environment with constant temperature of 25°C, constant humidity and 12 hr-12 hr light-dark cycle. Fly stocks used in this study are listed in Supplementary Table 5.

### Application of various stresses and stress removal

All *Oregon-R* (wild type) flies were well-fed and given additional yeast paste prior to application of stress. Ovaries of flies that were fed normal food at 25°C were used as controls. Flies were kept at 25°C unless stated otherwise. Heat shock was induced by keeping flies at 37°C for one hour. Oxidative stress was induced by keeping the flies in vials containing a filter paper soaked in solution of 0.05 M sodium arsenite and 5% sucrose for 3-3.5 hours. Starvation was induced by keeping the flies in vials containing a filter paper soaked in water (no sugar, no protein) for 24 hours. Heat shock stress was removed by keeping flies at 25°C for two hours. Oxidative stress was removed by keeping flies on 5% sucrose without sodium arsenite for three hours, and starvation was removed by keeping flies on normal food for 24 hours post-starvation.

### Immunohistochemistry

Ovaries were dissected in 1X cold phosphate buffered saline (PBS) in a dissection dish. Two forceps were used to remove the posterior-most cuticle. The abdomen was then gently squeezed in an anterior to posterior direction to retrieve the ovaries. Dissected ovaries were fixed in 4% paraformaldehyde (Polysciences Inc.) in PBT (pH 7.4) for 15 minutes on a rotator at highest speed. Fixed ovaries were washed in PBT twice quickly, followed by 2 washes on nutator for 15 minutes each. Samples were blocked in PBTB 2 times for 15 minutes each. Samples were nutated overnight at 4°C in appropriate dilution of primary antibody (diluted in PBTB). Ovaries were washed twice quickly, followed by 2 times washes for 15 minutes in PBT. They were blocked in PBTB for 30 minutes. Appropriate secondary antibodies were added (1: 500 pre-diluted in PBTB) and nutated for 2.5 hours at room temperature. Samples were washed 3 times for 15 minutes each. DAPI was added to the next PBT wash and nutated for 30 minutes. A final wash was performed using PBT for 5-10 minutes. The ovaries were stored in NPG + glycerol at 4°C overnight. For mounting, ovaries were separated into their individual ovarioles using syringe needles. All the buffers used are the same as stated in immunocytochemistry. List of primary and secondary antibodies used are listed in Supplementary Table 6. Fluorescence images were taken using a confocal laser-scanning microscope with airyscan (Zeiss LSM 900). The images were processed with Fiji (ImageJ), IMARIS, Adobe Photoshop and Adobe Illustrator.

### Counting of granules

Granules were counted using Cell Profiler 4.3.1. To count cytoplasmic AGO1 granules in a particular egg chamber, DAPI channel was used as a mask over the AGO1 channel, and the masked image was used to count the cytoplasmic granules of defined size range (particles having 8-15-pixel unit diameter). Total pixel area of the AGO1 channel and pixel area of only cytoplasmic AGO1 (masked image) was calculated using Cell profiler. Cytoplasmic AGO1 area in µm^2^ was calculated by multiplying pixel area of cytoplasmic AGO1 with total area of the original image in µm^2^ (taken from Zeiss LSM 700) and then dividing this by total pixel area of AGO1. Number of cytoplasmic AGO1 granules per µm^2^ were calculated. Number of granules in each condition were represented relative to the control.

### AGO1 and Fibrillarin colocalization analysis

Fluorescence intensity profile plots across a selected region of interest in the images co-immunostained with AGO1 and Fibrillarin were generated using ImageJ (Plugins → Graphics → RGB Profile Plot).

### Co-immunoprecipitation and western blot analysis in *Drosophila*

Ovarian lysates for general co-immunoprecipitation (co-IP) were prepared from approximately 1-week-old *AGO1::GFP* flies that were homogenized with VWR Disposable Pellet Mixers and lysed in RIPA buffer containing 50 mM Tris-HCl (pH 7.5), 125 mM NaCl, 5% glycerol, 0.5% NP40, 0.25% Na-deoxycholate, 1.5 mM MgCl2, 1 mM dithiothreitol, 25 mM NaF, 1 mM Na3VO4, 1 mM EDTA, 2 mM EGTA, and protease inhibitors. Samples were then centrifuged at 15,000 *g* for 15 min at 4 °C, and then 1.3 mg of supernatants was immunoprecipitated with GFP-Trap beads (ChromoTek) following the manufacturer’s instructions. The presence of immunocomplexes was analyzed via SDS-PAGE and western blotting as described in (97). The following antibodies were used: mouse anti-Fibrillarin (1:1000,), mouse anti-GFP (1:500, Santa Cruz Biotechnology, sc-9996).

### snoRNA Sequences and rRNA Targets

The sequences of box C/D snoRNAs Me18S-G1358a, Me18S-G1358b, and Me28S-A3407a were obtained from *Drosophila melanogaster* genomic annotations. Complementary sequences to 18S and 28S rRNAs were identified by aligning candidate guide regions (between C and D boxes) with rRNA sequences from FlyBase (release r6.50).

### Clonal AGO1 loss-of-function experiment

Homozygous AGO1 loss-of-function follicular epithelium cell clones were generated using heat shock inducible Flp/FRT system. We crossed *yw; FRT 42B AGO1^LOF^/Cyo* to *hsFlp; FRT 42B GFP/Cyo.* This *AGO1^LOF^* allele has a C-T mutation. Due to this mutation, amino acid 212 Q is changed to a stop codon in AGO1 isoform B, and amino acid 246 Q is changed to a stop codon in AGO1 isoform A and C. This produces AGO1 protein that lacks both important domains, PAZ and PIWI. To induce *AGO1 ^LOF^* follicular epithelium cell clones, adult progeny from the above cross were heat shocked for one hour in a 37°C water bath in empty vial for two consecutive days. After 3-4 days, ovaries were dissected and subsequently analyzed using immunohistochemistry. Cells lacking GFP and expressing AGO1 loss-of-function allele (*hsFlp; FRT42B AGO1^LOF^*) were compared with neighboring heterozygous control cells (*hsFlp; FRT42B GFP/FRT42B AGO1^LOF^*, GFP-positive).

### Clonal RNAi and overexpression experiments

The heat shock-inducible hsFlp; act>CD2>Gal4, UAS-GFP system also called the Flp-out method, was used to generate various RNAi and overexpression follicular epithelium cell clones. Myc clonal RNAi and overexpression experiments were performed by crossing *hsFlp; act>FRT-CD2-FRT>Gal4, UAS-GFP* CD2 to *UAS-Myc^RNAi^* (BDSC # 36123) and crossing *hsFlp; act>FRT-CD2-FRT>Gal4, UAS-GFP to w; UAS-Myc* (BDSC # 9674), respectively. AGO1 clonal RNAi experiment was conducted by crossing *hsFlp; act>FRT-CD2-FRT>Gal4, UAS-GFP* to *UAS-AGO1^RNAi^* (BDSC # 33727)*. Dcr1* clonal RNAi experiments were conducted by crossing *hsFlp; act>FRT-CD2-FRT>Gal4, UAS-GFP* to *eyFlp; +; FRT 82B UAS-dcr^RNAi^ D482/TM3* (98). Adult progeny from these crosses were heat shocked for one hour in a 37°C water bath in empty vial for two consecutive days. After 3-4 days, ovaries were dissected and subsequently analyzed using immunohistochemistry. After heat shock induction, GFP-positive clonal cells with downregulated or upregulated levels of the protein of interest are surrounded by control GFP-negative cells, which allows their comparison in the same egg chamber.

### Measurement of cell and nucleolar area

Area of the selected clonal and surrounding control region (selected using cadherin staining which marks cell boundaries) was measured using ImageJ (using Analyze → Measure). Then, the area was divided by the number of cells present in the region to obtain area per cell. Nucleolar area was obtained by measuring area enclosed by Fibrillarin staining (ImageJ using Analyze → Measure). Mean cell area and mean nucleolar area were calculated and represented as Mean±SEM relative to the control. Student’s t-test was applied for statistical analyses.

### RNA extraction and reverse transcription

Ovaries were homogenized in 200µl of Trizol reagent (Ambion). 50µl of chloroform was added to the homogenized samples. Tubes were shaken vigorously by hand for 15 seconds and incubated them at 15 to 30°C for 2 to 3 minutes. Samples were centrifuged at 12,000× g (max) for 15 minutes at 2 to 8°C. The top aqueous phase overlying the interphase was transferred to a new tube and 0.1ml of 100% isopropanol was added to it. Samples were stored at 15 to 30°C for 10 min and centrifuged at 12000 x g for 10 minutes. The supernatant was removed and the pellet was washed with 75% Ethanol by vortexing and centrifuged at 7500 x g for 5 min. The pellet was air dried and dissolved in DEPC treated water (20-50µl). Up to 4µg of RNA was treated with DNase (Ambion) in 40µl reaction at 37°C for 40 minutes. EDTA was added to the reaction (5mM final concentration of EDTA desired) followed by deactivation of DNase at 75°C for 10 minutes. 1µg of DNase treated RNA was reverse transcribed using random primers with High Capacity Reverse Transcriptase (Applied Biosystems) in a 20µl reaction. Reaction was set up according to the manufacturer’s instructions.

### Quantitative-PCR (qPCR)

The qPCR was performed using Fast SYBR Green reagents (Qiagen) in Step One Plus Real Time PCR System (Applied Biosystems) according to the manufacturer’s instructions. Each reaction was set up with 300nM of forward and reverse primer concentrations and 20ng cDNA as template in 20μl volume. Primers for AGO1 (forward GAGGTGCTGGACATTCGCG and reverse GACGCATCTGGCCGCAGT) and pre-rRNA (forward GTTAGTGTGGGGCTTGGCAAC and reverse CGCCGTTGTTGTAAGTACTCGC) were designed manually (NEB Tm calculator was used). All other primers sequences were obtained from DRSC FlyPrimerBank (https://www.flyrnai.org/flyprimerbank). All primers were ordered from Microsynth. The analysis of the acquired C_T_ values was done using StepOne Software (Applied Biosystems) and Microsoft Excel. ΔC_T_ values were calculated by subtracting the average C_T_ value of the technical replicates of the respective genes from average C_T_ values of housekeeping gene, *α-tubulin* (*α-Tubulin 84B* forward CACACCACCCTGGAGCATTC and reverse CCAATCAGACGGTTCAGGTTG). ΔΔC_T_ was calculated by subtracting ΔC_T_ of the treated/mutant sample from ΔC_T_ of the control sample. Finally, the ΔΔC_T_ values were used to calculate the relative gene expression levels using the formula: 2^-ΔΔC^_T_. Two-tailed Student’s t-test was used for the calculation of p-values for statistical analyses.

### Processing and analysis of AGO-bound small RNAseq data

Publicly available AGO-bound small RNAseq data was used for our analysis (38). Raw data from their study was deposited in Gene Expression Omnibus (GEO) with accession number GSE129600. We downloaded FASTQ files of three biological replicates of *Drosophila* S2 cell samples (*Control*) and *Drosophila* S2 cells overexpressing AGO1 (*AGO1^OE^*) (sample numbers - GSM3733652, GSM3733653, GSM3733654, GSM3733657, GSM3733658, GSM3733659). These files were imported to StrandNGS software (version 3.4) to map sequencing reads with the *Drosophila* genome. The normalization of reads was performed using DESeq algorithm in StrandNGS. Average of three biological replicates was used to find the percentage of each type of small RNA detected and most abundant snoRNAs. Differential snoRNA expression analysis between *AGO1^OE^ and control* was performed using R statistical software. gplots and RColorBrewer packages were used for data visualization.

### Protein sequences and domain annotation

Drosophila protein sequences were obtained from FlyBase (https://flybase.org). AGO protein domains were annotated using Simple Modular Architecture Research Tool (SMART) (http://smart.embl-heidelberg.de). and were subject to further manual annotations based on evolutionary analysis with known human AGO sequences using ClustalW, Jalview, and comparative structural mapping using PyMOL. The results of protein domain mapping as well as the prediction outcome was visualized and analyzed using the PyMOL Molecular Graphics System version 3.0.5 (Schrödinger, LLC) and ChimeraX (UCSF) (99).

For the purpose of multiple sequence alignment, resources of AGO1 amino acid sequence were (1) FlyBase: Drosophila melanogaster (Q7KY08; Q7KY08_DROME; FBpp0086740; AGO1-RC) (2) UniProt-SWISSPROT: Mus musculus (Q8CJG1; AGO1_MOUSE); Homo sapiens (Q9UL18; AGO1_HUMAN) (3) UniProt-TREMBL: Mayetiola destructor (K7Y0I3; K7Y0I3_MAYDE); Danio rerio (K4I6K9;K4I6K9_DANRE); Xenopus tropicalis (A0A6I8QDR; A0A6I8QDR2_XENTR); Crocodylus porosus (A0A7M4FM46; A0A7M4FM46_CROPO); Gallus gallus (A0A1D5PXD8; A0A1D5PXD8_CHICK); Gorilla gorilla gorilla (G3S0G5; G3S0G5_GORGO). Regarding AGO-1, the well annotated UniProt database contained the high confidence AGO1 entries of mouse and human (each with 857 AA). FlyBase contained 4 entries of AGO1, with three identical peptide sequences of 984 AA with extended aminoterminal sequence, and one shorter isoform of 950 AA that we chose and that still displayed an aminoterminal extension when compared with human and mouse sequences. Further representative sequences of further species were selected from UniProt-Trembl with focus on similar sizes as for the respective UniProt.Swissprot entries (857 AA). We also selected a further species of the order of diptera. Of note, UniProt-SWISSPROT contains even a high-confident entry of AGO1 of Schizosaccharomyces pombe, with a high degree of similarity when compared to higher eukaryotes and with the alternative name “Cell cycle control protein ago1”. It is annotated as “Required for G1 arrest and mating in response to nitrogen starvation.” (Sequence not integrated in MSA due to still representing an outlier).

### Prediction of ribonucleoprotein complexes

Initial complexes comprising AGO1, the Fibrillarin-NOP56/58-SNU13 assembly, and RNAs (box C/D snoRNA Me18S-G1359a, also referred to as CD39b, and a fragment of its 18S rRNA target encompassing the methylation site) were modeled using AlphaFold3 (77). The resulting AGO1-RNA and Fibrillarin-NOP56/58-SNU13-RNA complexes were physically plausible, whereas models containing both AGO1 and the complete Fibrillarin-NOP56/58-SNU13 assembly lacked functionally relevant interactions between the protein and RNA components.

To obtain a model of the complete AGO1-Fibrillarin-NOP56/58-SNU13-RNA complex, initial unbiased protein-protein docking of the AGO1 and Fibrillarin subunits, without the remaining protein and RNA components, was performed using HDOCK (79). The resulting AGO1-Fibrillarin docking poses were evaluated in the context of the complete ribonucleoprotein assembly by reconstructing the remaining components from the AlphaFold3-generated subcomplexes. At this stage, the snoRNA was represented as two disconnected fragments corresponding to the regions bound to AGO1 and the Fibrillarin-containing snoRNP, respectively. Docking poses were assessed for whether the relative orientation of AGO1 and Fibrillarin would permit the two snoRNA fragments to be connected by the intervening RNA segment, while avoiding major steric clashes between AGO1 and the other components of the Fibrillarin-NOP56/58-SNU13 assembly. Poses incompatible with these spatial requirements were discarded, and the remaining models were used to identify a plausible AGO1-Fibrillarin interaction interface, including residues 218 and 220 of the Fibrillarin domain and residues 472 and 475 of AGO1.

The AGO1-Fibrillarin interface was subsequently refined by restraint-guided protein-protein docking using HADDOCK (80), using the above-mentioned residues of Fibrillarin and AGO1 as active residues for ambiguous interaction restraints. The resulting AGO1-Fibrillarin models were used to reconstruct a complete AGO1-Fibrillarin-NOP56/58-SNU13-RNA assembly.

The model was first refined using SimRNA (81), to restore the continuity of the snoRNA chain, bound both to AGO1 and the Fibrillarin subcomplex. At this stage, only residues 6–18 (broken linker) and 58–74 (3’ terminus) of the snoRNA were allowed to be fully flexible, while the remaining parts of the snoRNA and the substrate fragment were frozen. A best-scoring model with the optimized RNA chain was identified from a SimRNA simulation run with default settings for 1,000,000 Monte Carlo steps. That model subsequently underwent energy minimization in GROMACS 2021.2. The AMBER99SB-ILDN and AMBER94 force fields (100) were used for the protein and RNA components, respectively.

Models were visualized and analyzed using PyMOL version 3.0.5 (Schrödinger, LLC) and ChimeraX (UCSF) (99). RNA-binding modes were evaluated by comparison with experimentally determined RNA-protein structures. Predicted Fibrillarin-RNA interactions were interpreted in the context of archaeal and eukaryotic box C/D RNP complexes, whereas AGO1-RNA interactions were assessed against available AGO-RNA structures.

