## Supplementary material for "Argonaute 1 associates with nucleolar snoRNP machinery and regulates pre-rRNA levels and cell growth": Supplmentary Data

### SUPPLEMENTARY DATA

#### Supplementary Figures and Legends

##### Supplementary Figure 1

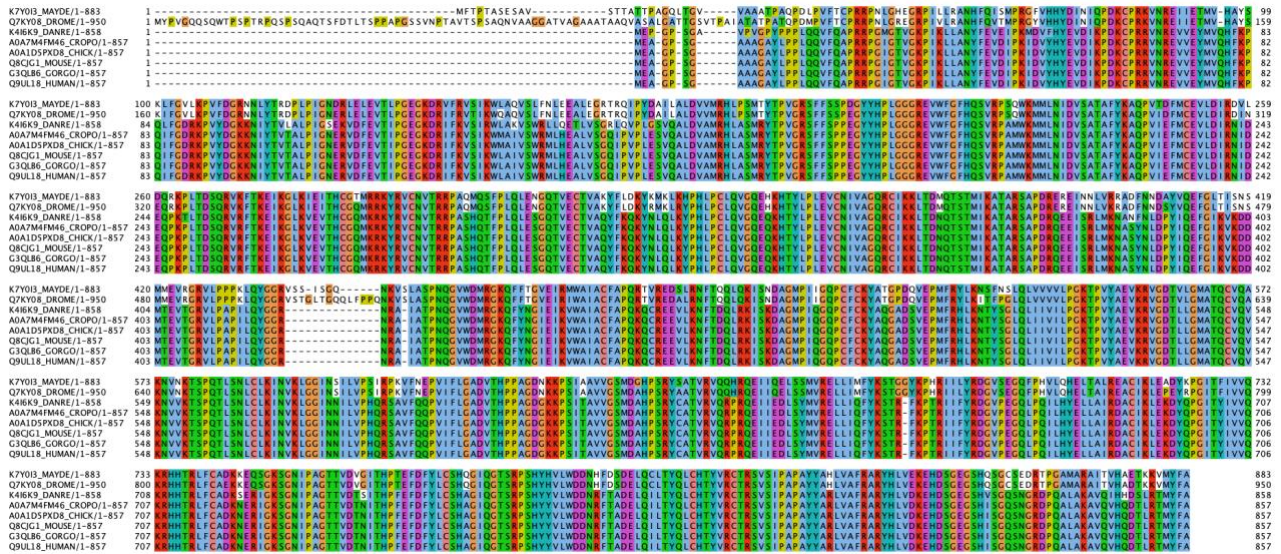

##### Supplementary Figure 1. Multiple sequence alignment of AGO1 protein sequences

The multiple sequence alignment of several eukaryote protein sequences reveals a huge degree of sequence conservation. Differences between the proteins arise largely from the dAGO1 N-terminal region, which is elongated when compared to hAGO1, a typical feature of *Drosophila* protein sequences. Without this part the sequence identity between dAGO1 and hAGO1 is 73%. Sequences were extracted from FlyBase and UniProt (K7Y0I3\_MAYDE: Mayetiola destructor; Q7KY08\_DROME: *Drosophila melanogaster*; K4I6K9\_DANRE: *Danio rerio*; A0A7M4FM46\_CROPO: *Crocodylus porosus*; A0A1D5PXD8\_CHICK: *Gallus gallus*; AGO1\_MOUSE: *Mus musculus*; G3SQB6\_GORGO: *Gorilla gorilla*; AGO1\_HUMAN: *Homo sapiens*).

Supplementary Figure 2

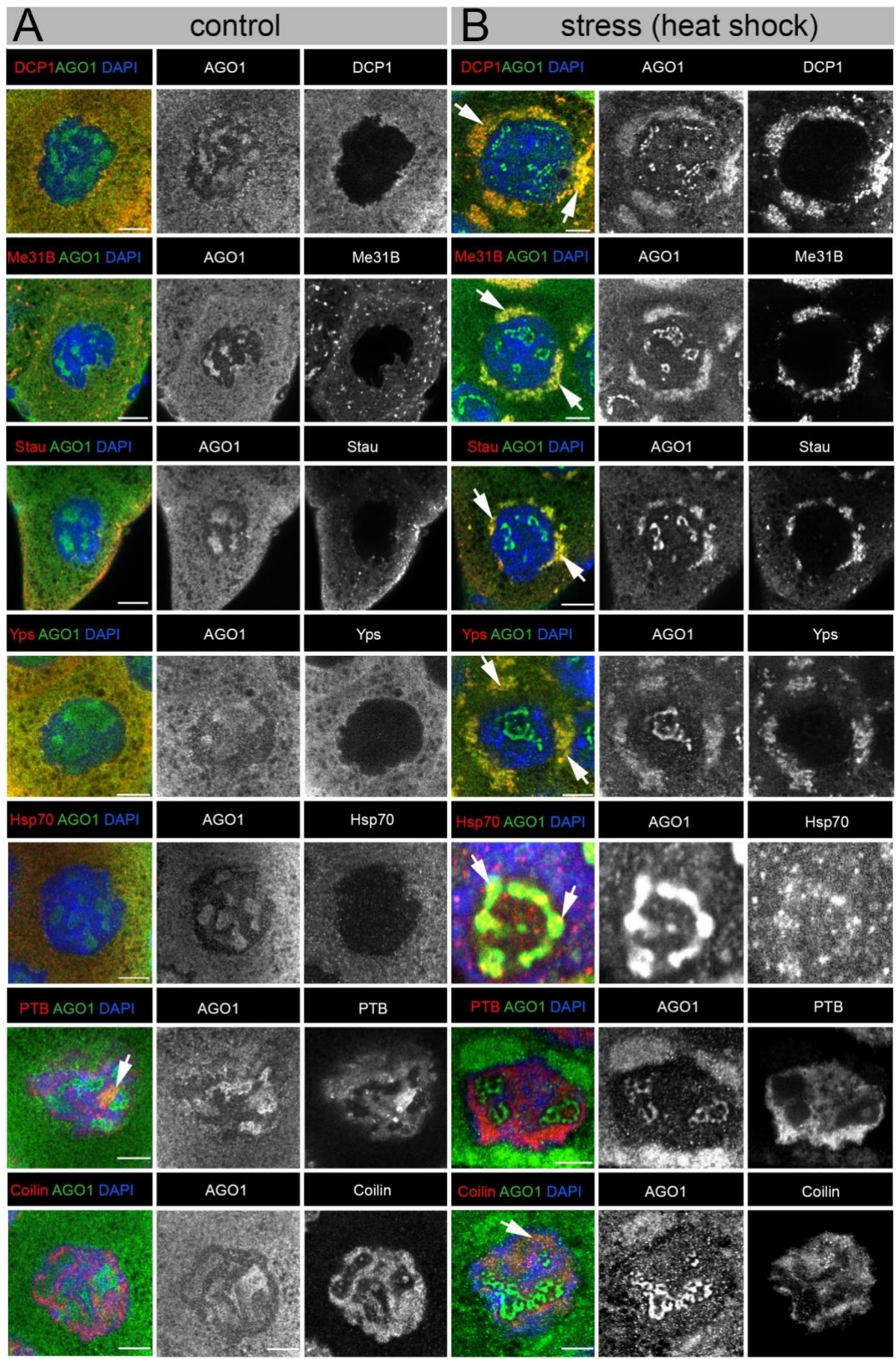

#### **Supplementary Figure 2. AGO1 colocalizes with known RNP granules in the cytoplasm and in the nucleus**

**A-B.** Images show nurse cells (st.10) stained with anti-AGO1 (green) and anti-RNP marker (red) and if AGO1 colocalizes with a given RNP marker (yellow). DCP1, Me31B, Staufen (Stau) Yps and Hsp70 are cytoplasmic RNP markers. PTB and Coilin are nuclear RNP markers. Anti-GFP antibody is used for staining DCP1-GFP, Me31B-GFP, Staufen-GFP, Yps-GFP and PTB-GFP. Anti-Coilin antibody is used to mark Cajal bodies, anti-Hsp70 marks stress granules.

Under control conditions (**A**), AGO1 (green) has a dispersed pattern in the cytoplasm and a clear distinct pattern in the nucleus (A). Upon heat shock, AGO1 forms a cloud-like pattern in the cytoplasm around the nuclear periphery, and its nuclear pattern becomes more compact (**B**).

Under control conditions, DCP1 distribution pattern is dispersed in the cytoplasm and weak colocalization is seen with AGO1. B. Upon heat shock, DCP1 accumulates around the nucleus in the same cloud-like pattern as AGO1 (white arrows) and strongly colocalizes with it (yellow).

Granules of Me31B are observed in control in the cytoplasm and these granules colocalize with AGO1 (white arrows). Strong colocalization of Me31B with AGO1 is observed surrounding the nuclear region upon heat shock (yellow).

Under control conditions, a few granules of Staufen are observed especially around the edges of the egg chamber, where it weakly colocalizes with AGO1 (white arrows). Upon heat shock, Staufen strongly colocalizes with AGO1 surrounding the nucleus.

Yps has a dispersed pattern in the cytoplasm under control conditions and weakly colocalizes with AGO1. Yps strongly colocalizes with AGO1 in the same cloud-like pattern (yellow) upon heat shock.

Under control conditions, AGO1 shows weak cytoplasmic colocalization with Hsp70, which is absent from the nucleus. Following heat shock, Hsp70 accumulates in the nucleus, where it strongly colocalizes with AGO1.

PTB is predominantly present in the nucleus and colocalizes with AGO1 under control conditions (white arrow). Upon heat shock, the nuclear localization pattern of both PTB and AGO1 is altered, and their colocalization is no longer detected.

Under control conditions, AGO1 does not colocalize with Coilin. Upon heat shock, AGO1 colocalizes with Coilin in distinct granules.

All images are single Z-sections. Scale bars = 10µm.

Supplementary Figure 3

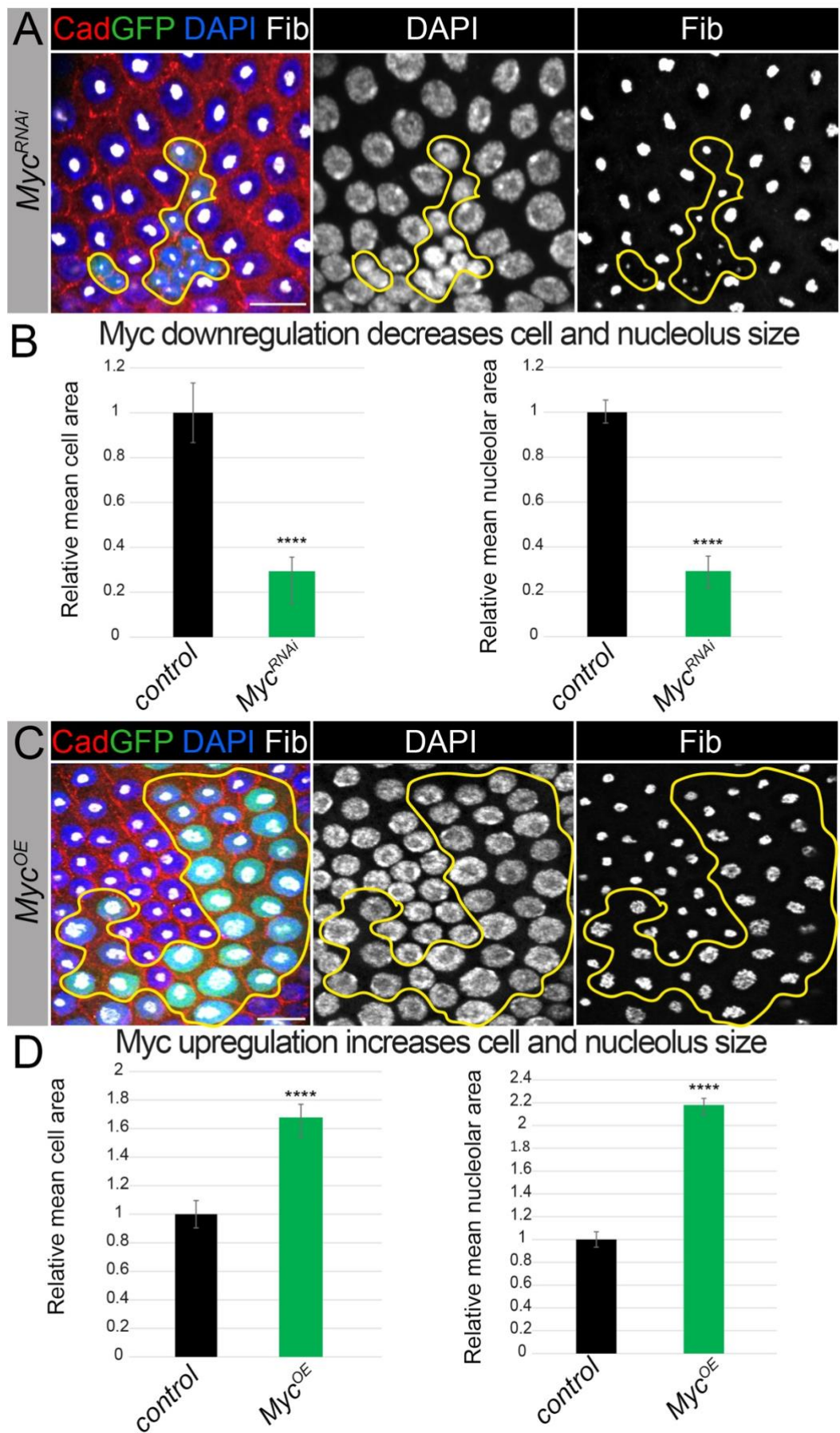

##### **Supplementary Figure 3. Myc downregulation and overexpression affects cell and nucleolar size**

**A.** Follicular epithelium cell clones with downregulated Myc (*act>Myc<sup>RNAi</sup>, GFP*, outlined by yellow line) have much smaller cells and nucleoli than the neighboring control cells.

**B.** Mean cell and nucleolar areas are significantly decreased in *Myc<sup>RNAi</sup>* clones compared to the respective means of the neighboring control cells (n= 13 clones and n= 36 cells from 13 clones).

**C.** Follicular epithelium cell clone overexpressing Myc (*act>Myc<sup>OE</sup>, GFP*, outlined by yellow line) has larger cells and (D'') nucleoli than the neighboring control cells.

**D.** Mean cell and nucleolar areas are significantly increased in *Myc<sup>OE</sup>* clones compared to the respective means of the neighboring control cells (n= 11 clones and n=32 cells from 11 clones).

All images are maximum intensity projections of multiple Z-sections. DAPI marks the nuclei (blue). Fibrillarin (Fib) marks the nucleoli (grey). Cadherin (Cad) marks the cell boundaries (red). Scale bars = 10µm. Cell and nucleolar area were measured using ImageJ.

Mean cell area and mean nucleolar (Mean±SEM) are represented relative to the control. Student's t-test was applied for statistical analyses, \*\*\*\*p-value <0.0001.

#### Supplementary Tables

**Supplementary Table 1. Summary of AGO1 colocalization with known RNP markers**

| RNP Marker | Symbol | Type of RNP granule | Location | Function | Colocalization with AGO1<br>Control Stress |  |
| --- | --- | --- | --- | --- | --- | --- |
| Decapping protein 1 | DCP1 | processing body, sponge body | cytoplasm | Decapping enzyme - functions in translational repression | + | ++ |
| Maternal expression at 31B | Me31B | processing body, nuage, sponge body | cytoplasm | Decapping activator/RNA helicase - functions in translational repression | + | ++ |
| Staufen | Stau | processing body | cytoplasm | RNA binding protein playing a role in proper mRNA localization | + | ++ |
| Ypsilon Schachtel | Yps | processing body | cytoplasm | Y-box protein regulating RNA localization and translation | + | ++ |
| Heat-shock-protein-70 | Hsp70 | stress granule | cytoplasm | Heat-shock protein Facilitates cell survival during stress | + | ++ |
| Polypyrimidine tract-binding protein | PTB | high-order RNP assembly | nucleus | Regulation of RNA processing including alternative splicing | + | - |
| Coilin | Coil | Cajal body | nucleus | Required for the formation of Cajal bodies | - | + |
| Fibrillarin | Fib | nucleolus | nucleus | rRNA methyltransferase involved in rRNA processing | ++ | ++ |

- no colocalization; + weak colocalization; ++ strong colocalization

**Supplementary Table 2. AGO-bound small RNA reads**

| Control |  |  | *AGO1<sup>OE</sup>* |  |  |
| --- | --- | --- | --- | --- | --- |
| small RNA type | No. of reads | % of total reads | small RNA type | No. of reads | % of total reads |
| *miRNAs* | 757.9853 | 32.39% | *miRNAs* | 701.3626 | 29.96% |
| *snoRNAs* | 190.5891 | 8.15% | *snoRNAs* | 219.3997 | 9.37% |
| *snRNAs* | 105.5636 | 4.51% | *snRNAs* | 116.1045 | 4.96% |
| *tRNAs* | 1286.338 | 54.97% | *tRNAs* | 1304.852 | 55.73% |
| **Total reads** | 2340.476 | 100% | **Total reads** | 2341.719 | 100% |

**Control** - *Drosophila* S2 cells; ***AGO1<sup>OE</sup>*** - *Drosophila* S2 cells overexpressing AGO1

tRNAs - transfer RNAs; miRNAs - microRNAs; snoRNAs - small nucleolar RNAs; snRNAs - small nuclear RNAs

\* - analyzed using publicly available data (1)

**Supplementary Table 3. Most abundant AGO-bound snoRNAs**

| snoRNA | Type of snoRNA | No. of snoRNA reads in control | % of total snoRNA reads in control |
| --- | --- | --- | --- |
| *snoRNA:Me18S-A1576* | C/D box | 5.89 | 3.09 |
| *snoRNA:Or-CD12* | orphan | 5.10 | 2.68 |
| *snoRNA:Me28S-G2703a* | C/D box | 4.63 | 2.43 |
| *snoRNA:MeU6-C68* | C/D box | 4.57 | 2.40 |
| *snoRNA:Me18S-A425* | C/D box | 4.18 | 2.19 |
| *snoRNA:Me28S-G2017* | C/D box | 4.16 | 2.18 |
| *snoRNA:U3:54Aa* | C/D box | 4.08 | 2.14 |
| *snoRNA:Me28S-C3227b* | C/D box | 4.03 | 2.12 |
| *snoRNA:Me18S-A1061* | C/D box | 3.92 | 2.06 |
| *snoRNA:U3:9B* | C/D box | 3.52 | 1.85 |
| *snoRNA:U3:54Ab* | C/D box | 3.47 | 1.82 |
| *snoRNA:Me28S-G1083a* | C/D box | 3.43 | 1.80 |
| *snoRNA:Me28S-C3227a* | C/D box | 3.33 | 1.75 |
| *snoRNA:Me28S-G1083c* | C/D box | 3.02 | 1.59 |
| *snoRNA:Me28S-G1083d* | C/D box | 2.87 | 1.50 |
| *snoRNA:Me18S-G1620* | C/D box | 2.71 | 1.42 |
| *snoRNA:Psi28S-1192a* | H/ACA box | 2.63 | 1.38 |
| *snoRNA:Me5.8S-G74* | C/D box | 2.57 | 1.35 |
| *snoRNA:Me28S-G980* | C/D box | 2.44 | 1.28 |
| *snoRNA:Me18S-A1806* | C/D box | 2.44 | 1.28 |
| *snoRNA:Or-CD2* | orphan | 2.42 | 1.27 |
| *snoRNA:Me28S-C2645c* | C/D box | 2.35 | 1.23 |
| *snoRNA:Me28S-C2645b* | C/D box | 2.35 | 1.23 |
| *snoRNA:Me28S-G1083b* | C/D box | 2.30 | 1.21 |
| *snoRNA:Me28S-A2589b* | C/D box | 2.28 | 1.19 |
| *snoRNA:Psi18S-1377a* | H/ACA box | 2.22 | 1.16 |
| *snoRNA:Me28S-G3113a* | C/D box | 2.20 | 1.15 |
| *snoRNA:Me28S-G2173* | C/D box | 2.09 | 1.10 |
| *snoRNA:Me28S-A982a* | C/D box | 2.01 | 1.05 |
| *snoRNA:Me28S-G3255a* | C/D box | 1.97 | 1.03 |

| snoRNA | Type of snoRNA | No. of reads in AGO1<sup>OE</sup> | % of total snoRNA reads in AGO1<sup>OE</sup> |
| --- | --- | --- | --- |
| **snoRNA:Me18S-A1576** | C/D box | 6.34 | 2.89 |
| **snoRNA:Me28S-G1083c** | C/D box | 5.71 | 2.60 |
| **snoRNA:Me28S-G2703a** | C/D box | 5.63 | 2.57 |
| **snoRNA:Me28S-G1083a** | C/D box | 5.62 | 2.56 |
| **snoRNA:MeU6-C68** | C/D box | 5.11 | 2.33 |
| **snoRNA:Or-CD12** | orphan | 4.96 | 2.26 |
| **snoRNA:U3:54Aa** | C/D box | 4.51 | 2.05 |
| **snoRNA:Psi28S-1192a** | H/ACA box | 4.28 | 1.95 |
| **snoRNA:Me28S-C3227b** | C/D box | 4.05 | 1.85 |
| **snoRNA:Me28S-G1083b** | C/D box | 3.93 | 1.79 |
| **snoRNA:Me18S-A425** | C/D box | 3.85 | 1.75 |
| **snoRNA:Me18S-G1620** | C/D box | 3.65 | 1.67 |
| **snoRNA:Me28S-G1083d** | C/D box | 3.63 | 1.65 |
| **snoRNA:Me28S-G3255a** | C/D box | 3.58 | 1.63 |
| **snoRNA:Or-CD2** | orphan | 3.41 | 1.55 |
| **snoRNA:U3:54Ab** | C/D box | 3.38 | 1.54 |
| **snoRNA:Me18S-G1358b** | C/D box | 3.26 | 1.49 |
| **snoRNA:Me28S-G2017** | C/D box | 3.19 | 1.45 |
| **snoRNA:Me18S-A1061** | C/D box | 3.14 | 1.43 |
| **snoRNA:Me28S-A982a** | C/D box | 3.07 | 1.40 |
| **snoRNA:Me18S-C1366** | C/D box | 2.99 | 1.36 |
| **snoRNA:U49:66Db** | C/D box | 2.98 | 1.36 |
| **snoRNA:U3:9B** | C/D box | 2.92 | 1.33 |
| **snoRNA:Me18S-C1831** | C/D box | 2.83 | 1.29 |
| **snoRNA:Me28S-A1322** | C/D box | 2.81 | 1.28 |
| **snoRNA:U76:54Eb** | C/D box | 2.72 | 1.24 |
| **snoRNA:Me18S-G1358a** | C/D box | 2.66 | 1.21 |
| **snoRNA:Me28S-C3227a** | C/D box | 2.65 | 1.21 |
| **snoRNA:Or-CD4** | orphan | 2.63 | 1.20 |
| **snoRNA:Me28S-A2589b** | C/D box | 2.34 | 1.07 |

Control - *Drosophila* S2 cells; **AGO1<sup>OE</sup>** - *Drosophila* S2 cells overexpressing AGO1
**snoRNAs** indicated in bold are common to both control and AGO1<sup>OE</sup>; other snoRNAs are non-overlapping between control and AGO1<sup>OE</sup>

\* -analyzed using publicly available data (1)

**Supplementary Table 4. List of differentially represented AGO-bound snoRNAs upon AGO1 overexpression \***

| **snoRNA** | **Log<sub>2</sub> fold change** | **Fold change upon AGO1 overexpression** |
| --- | --- | --- |
| **snoRNA:Me18S-G1358a** | 1.187742 | 2.27796 |
| **snoRNA:Me28S-A3407a** | 1.108933 | 2.156861 |
| **snoRNA:Psi18S-996** | 1.060561 | 2.085743 |
| **snoRNA:Me28S-C3351** | 1.048253 | 2.068024 |
| **snoRNA:Me28S-A771** | 0.967995 | 1.95612 |
| **snoRNA:Me28S-A2589c** | 0.894178 | 1.858551 |
| **snoRNA:Me28S-G3081a** | 0.795607 | 1.735807 |
| **snoRNA:Or-CD4** | 0.793729 | 1.733549 |
| **snoRNA:Psi18S-1854a** | 0.775169 | 1.711391 |
| **snoRNA:Me28S-A982b** | 0.772942 | 1.70875 |
| **snoRNA:Or-CD1** | 0.766185 | 1.700766 |
| **snoRNA:Or-CD11b** | 0.763615 | 1.697739 |
| **snoRNA:U14:30Ea** | 0.743137 | 1.673812 |
| **snoRNA:Me28S-G1083c** | 0.738338 | 1.668253 |
| **snoRNA:Me28S-A1322** | 0.728713 | 1.65716 |
| **snoRNA:Me18S-C1831** | 0.678532 | 1.60051 |
| **snoRNA:U76:54Eb** | 0.671828 | 1.59309 |
| **snoRNA:Psi28S-3316c** | 0.668216 | 1.589107 |
| **snoRNA:snR38:54Ec** | 0.650713 | 1.569944 |
| **snoRNA:Me28S-G3255a** | 0.624233 | 1.541391 |
| **snoRNA:Psi28S-3385a** | 0.614777 | 1.531321 |
| **snoRNA:MeU2-A31** | 0.609359 | 1.525581 |
| **snoRNA:Me28S-A2634c** | 0.598652 | 1.514301 |
| **snoRNA:Me28S-U3344a** | 0.596981 | 1.512548 |
| **snoRNA:Me18S-G1358b** | 0.585636 | 1.500701 |
| **snoRNA:MeU5-C46** | -0.62053 | -1.53744 |
| **snoRNA:Psi18S-1377a** | -0.6353 | -1.55326 |
| **snoRNA:Me28S-U1848** | -0.70167 | -1.62639 |
| **snoRNA:Psi18S-841a** | -0.7113 | -1.63728 |
| **snoRNA:Psi28S-3405a** | -0.74656 | -1.67778 |
| **snoRNA:Psi28S-3091b** | -0.89984 | -1.86585 |
| **snoRNA:Me28S-U2134a** | -1.01263 | -2.01759 |
| **snoRNA:Psi28S-2876** | -1.15776 | -2.2311 |

\* -analyzed using publicly available data (1)

**Supplementary Table 5. Key resources table**

| REAGENT or RESOURCE | SOURCE | IDENTIFIER |
| --- | --- | --- |
| **Antibodies** |  |  |
| Rabbit polyclonal anti-AGO1 (1:500) | Abcam | # ab5070 |
| Chicken anti-GFP (1:5000) | Aves Labs | # GFP-1020<br>Lot # GFP697986 |
| Guinea pig anti-Coilin (1:1000) | Joseph Gall |  |
| Mouse monoclonal anti-Fibrillarin (1:500) | Abcam | # ab4566 |
| Mouse monoclonal anti-LaminC (1:20) | DSHB | # LC28.26 |
| Rat monoclonal anti-Cadherin (1:20) | DSHB | # DCAD2 |
| Mouse monoclonal anti-HSP-70 (1:1000) | Enzo Life Sciences | # ADI-SPA-822-D,<br>RRID:AB_2039252 |
| Goat Alexa 568 anti-mouse (1:500) | Invitrogen | # A-11004 |
| Goat Alexa 568 anti-rabbit (1:500) | Invitrogen | # A-11011 |
| Goat Alexa 488 anti-mouse (1:500) | Invitrogen | # A-11001 |
| Goat Alexa 488 anti-rabbit (1:500) | Invitrogen | # A-11008 |
| Goat Alexa 488 anti-chicken (1:500) | Invitrogen | # A-11039 |
| Goat Alexa 568 anti-guinea pig (1:500) | Invitrogen | # A-11075 |
| Goat Alexa 647 anti-rat (1:500) | Invitrogen | # A-21247 |
| **Chemicals, peptides, and recombinant proteins** |  |  |
| Sodium arsenite | Sigma-Aldrich | S7400;CAS.# 7784-46-5 |
| **Critical commercial assays** |  |  |
| High-Capacity Reverse Transcriptase | Applied Biosystems | # 4368813 |
| Fast SYBR Green reagents | Qiagen | # 4385612 |
| **Experimental models: Organisms/strains** |  |  |
| *OregonR* | BDSC | # 5 |
| ***Dcp1-GFP N1*** | Brenneke Lab |  |
| ***EGFP Me31B/FM7; Sco/CyO*** | Ephrussi Lab |  |
| ***hsFlp; Stau-GFP; act>FRT-CD2-FRT>Gal4, UAS-GFP*** | Wu-Min Deng Lab |  |
| ***w; BDSC /CyO; Yps-GFP /TM6*** | Ephrussi Lab |  |
| ***mat-GFP PTB-wt/FM6*** | Ephrussi Lab |  |
| ***y[1] sc[\*] v[1] sev[21]; P[y[+t7.7] v[+t1.8]=TRiP.HMS01538]attP2*** | BDSC | # 36123 |
| ***w[1118]; P[w[+mC]=UAS-Myc.Z]132*** | BDSC | # 9674 |
| ***yw; FRT 42B AGO1LOF/Cyo*** | Carthew Lab |  |
| ***hsFlp; FRT 42B GFP/CyO*** | Shcherbata Lab |  |
| ***AGO::GFP*** | BDSC | # 50805 |
| ***y[1] sc[\*] v[1] sev[21]; P[y[+t7.7] v[+t1.8]=TRiP.HMS00610]attP2*** | BDSC | # 33727 |
| ***eyFlp;+;FRT 82B UAS-dcr<sup>RNAi</sup> D482/TM3*** | Carthew Lab |  |
| **Software and algorithms** |  |  |
| **ImageJ** | (2) | https://imagej.nih.gov/ij/ |
| **AlphaFold 2 and AlphaFold 3** | (3,4) | https://colab.research.google.com/github/sokrypton/ColabFold/blob/main/AlphaFold2.ipynb |
| **IMARIS** | Oxford Instruments, Zurich, Switzerland | https://www.oxinst.com |
| **CellProfiler** | (5) | https://cellprofiler.org |
